# Heparin nucleates and promotes tropoelastin coacervation through a transient interaction driven by domain 36

**DOI:** 10.64898/2026.04.30.721972

**Authors:** Robert Lu, Sean Reichheld, Min Jin, Simon Sharpe

## Abstract

Elastin is the extracellular matrix (ECM) protein responsible for the elastic recoil property of certain tissues including the skin, arteries, and lung. Elastic fibre assembly begins with the coacervation of soluble monomeric tropoelastin and is driven by interactions with other ECM components such as glycosaminoglycans (GAGs). Previous research shows that GAG interactions can promote tropoelastin coacervation but lack structural and mechanistic details of this interaction. In this study, we describe the key interactions between tropoelastin and heparin using NMR spectroscopy and coacervation experiments. We propose a mechanism in which substoichiometric GAGs can act as a nucleating scaffold, primarily through transient multivalent interactions with domain 36 of tropoelastin, reducing the energetic barrier for coacervation. Our results provide the first detailed molecular view of tropoelastin-GAG interactions and support a role for negatively charged GAGs in modulating tropoelastin coacervation and thus initiating elastic fibre assembly.

## Introduction

The extracellular matrix (ECM) is a complex, dynamic, and highly organized environment that mediates key biological functions, including cell adhesion, signalling, and the mechanical resilience of tissues [1]. Among ECM components, elastic fibres provide the elasticity required for the repeated deformation of tissues, such as the lung, skin, and arteries [2,3]. Elastic fibres consist of an elastin core surrounded by a fibrillin-rich microfibrillar matrix, and can vary greatly in morphology, elastin content [4], and mechanical properties [5,6] for different tissues. Most elastin synthesis occurs perinatally, during pregnancy and up to a year after birth [4,7,8], whereas in adulthood, elastin is extremely durable and has very little turnover and expression [9,10] – only in specific instances can elastin synthesis be reactivated, such as in wound repair and/or disease [4,11–13]. Thus, proper initial assembly of the elastic fibre is key to maintaining the structural integrity of tissues throughout life. Although the major steps of elastic fibre formation are understood at a macroscopic level, the molecular processes that regulate early tropoelastin assembly remain poorly defined. In particular, the molecular details of how specific ECM components, such as glycosaminoglycans (GAGs), interact with tropoelastin to modulate or regulate assembly remain unclear.

Tropoelastin is the ∼60 kDa monomeric precursor to elastin and is composed of alternating hydrophobic domains (HDs) and crosslinking domains (CLDs) [14]. Hydrophobic domains (HDs) of tropoelastin are low-complexity, repetitive and pseudorepetitive sequences consisting of mostly glycine, proline, and valine [15,16]. These regions are responsible for tropoelastin’s intrinsic disorder. The regular spacing of PG and VG motifs prevents the formation of extended secondary structure by favoring transient bends and turns [17–19]. HDs also mediate tropoelastin self-association through a phase separation event commonly referred to as coacervation [20–25]. Coacervation produces a dense yet hydrated and fluid protein-rich phase and a sparsely populated protein-poor phase [18,26–28]. Protein disorder is largely maintained in the dense phase because the process is driven by transient, multivalent, and non-specific interactions between hydrophobic residues [18,19,29–32]. Concentration of tropoelastin into a protein-rich dense phase is considered a critical step in organizing it for subsequent cross-linking by lysyl oxidase.

In contrast, crosslinking domains (CLDs) of tropoelastin contain 2-3 lysine residues spaced 2-3 residues apart within either an alanine-rich sequence (KA crosslinking domains) or a HD-like proline-rich sequence (KP domains) [14,15]. CLD lysine residues are involved in inter- and intramolecular crosslinks, which convert soluble tropoelastin monomers into the insoluble elastic fibre [33–35]. Positively charged residues are also found within the tropoelastin C-terminal domain 36, although they do not appear to contribute to crosslink formation and are not present in mature tissues [35,36]. Instead, domain 36, which contains an internal disulfide bond and the terminal charge cluster RKRK [15], has been proposed to play a role in facilitating ECM interactions that influence elastic fibre assembly [37,38].

Tropoelastin assembly in the ECM involves three main steps: coacervation, crosslinking, and deposition onto the microfibril. After secretion into the ECM by elastogenic cells such as smooth muscle cells, fibroblasts, and endothelial cells [2,4,11,39], tropoelastin coacervation is triggered by interactions with cell-surface GAGs [40–42]. The coacervate droplets gradually grow on the cell surface [25,43,44] while modification of lysine sidechains by lysyl oxidase results in formation of multifunctional crosslinks between tropoelastin crosslinking domains [4,34,35,45]. After initial coacervation and crosslinking, tropoelastin coacervates are deposited onto the fibrillin microfibril [24,38,46–52]. Current models of elastic fibre assembly have proposed that interactions between tropoelastin and lysyl oxidase or fibrillin are mediated by fibulins and matrix-associated glycoproteins (MAGPs) [49,53–67]. Once associated with the microfibril scaffold, further crosslinking and maturation stabilize the tropoelastin coacervate to produce the mature, insoluble elastic fiber [4,19,24,68,69].

While many ECM interactions are involved in controlling elastic fibre assembly, interactions with cell-surface GAGs may be particularly important as the first step in triggering tropoelastin coacervation. Previous research shows that the tropoelastin-GAG interaction can have varied effects, such as promoting tropoelastin coacervation [40–42,70–72], tropoelastin fibril formation [70,73,74], and cell adhesion [37,75,76], yet the underlying mechanism remains unclear. Some studies support a charged mode of interaction but do not identify the specific residues involved [42,71,77]. Others propose interactions with specific charged domains of tropoelastin, such as domains 15, 17, and 36, [37,75,76] showing how these domains can trigger cell spreading; however, whether these same interactions drive coacervation is uncertain. Additional studies suggest that non-charged interactions also occur, as GAGs can affect coacervation and fibril formation of elastin-like polypeptides (ELPs) solely composed of hydrophobic sequences [70,73]. Structural information upon GAG binding is limited. GAG binding appears to result in increased helicity of tropoelastin as measured by circular dichroism [41,78], but the specific domains involved are unknown. Consistent with a charge-mediated interaction, heparin tends to have a stronger effect in promoting tropoelastin coacervation than other GAGs such as chondroitin sulfate B and heparan [37,40,42], likely due to its higher degree of sulfation and resulting charge density [79]. In summary, although prior research supports an important role for GAGs in the initial stages of elastic fibre assembly, the molecular details of their interactions with tropoelastin and how they promote coacervation remain unresolved.

In this study, we investigate the molecular details of the interaction between tropoelastin and heparin, used here as an analog for the cell-surface GAG heparan sulfate. Using coacervation assays in conjunction with solution NMR, we identify the specific domains and residues of tropoelastin that mediate this interaction and determine how heparin influences early assembly events. Our data support a transient, charge-mediated interaction with heparin centered around the RKRK motif of domain 36 that subsequently allows heparin to interact with internal CLDs more readily. Heparin acts as a nucleating agent, binding multiple tropoelastin molecules, bringing them into closer proximity and reducing the energetic barrier for coacervation. These findings reveal the molecular mechanism by which GAGs promote tropoelastin coacervation and suggest that GAG type and abundance in the ECM may regulate the initiation of elastic fibre assembly.

## Results

### Construct selection and design

To investigate the molecular details of the interaction of heparin and tropoelastin and the effects on tropoelastin coacervation, we used smaller constructs representing fragments of tropoelastin as well as the full length human tropoelastin (hTE) (Figure 1, SI Tables S1, S2). These include fragments hTE2-14, which contains mostly KP crosslinking domains; hTE13-19, which contain proposed heparin binding sites within the central domains of tropoelastin [75,76]; hTE18-26, which contains the “hinge” region of tropoelastin with two adjacent KA crosslinking domains [80,81]; and hTE24-36, which includes both the proline-poor domain 30 and the positively charged domain 36, with the latter being identified as a heparin binding site [37,52]. These fragments accurately recapitulate local tropoelastin sequence propensities [82] and allow for the investigation of the role of different regions of tropoelastin. To facilitate investigation of specific sequences (here, domain 36) within a simplified background, our lab previously designed a 13 kDa elastin-like polypeptide (ELP) called ELP_3_, with similar alternating domain organization as tropoelastin, and sharing the ability to both coacervate and form crosslinked materials [19]. The ELP constructs consist of representative hydrophobic and crosslinking domain sequences derived from amniotic tropoelastin sequences [83–85], with ELP_3_(36) containing tropoelastin domain 36 in place of a C-terminal crosslinking domain. ELP29-36 consists of tropoelastin domains 29 – 36 with an ELP hydrophobic and crosslinking domain added to the N-terminus to make its molecular weight comparable to ELP_3_(36) while containing the natural C-terminal domains of tropoelastin.

**Figure 1.**
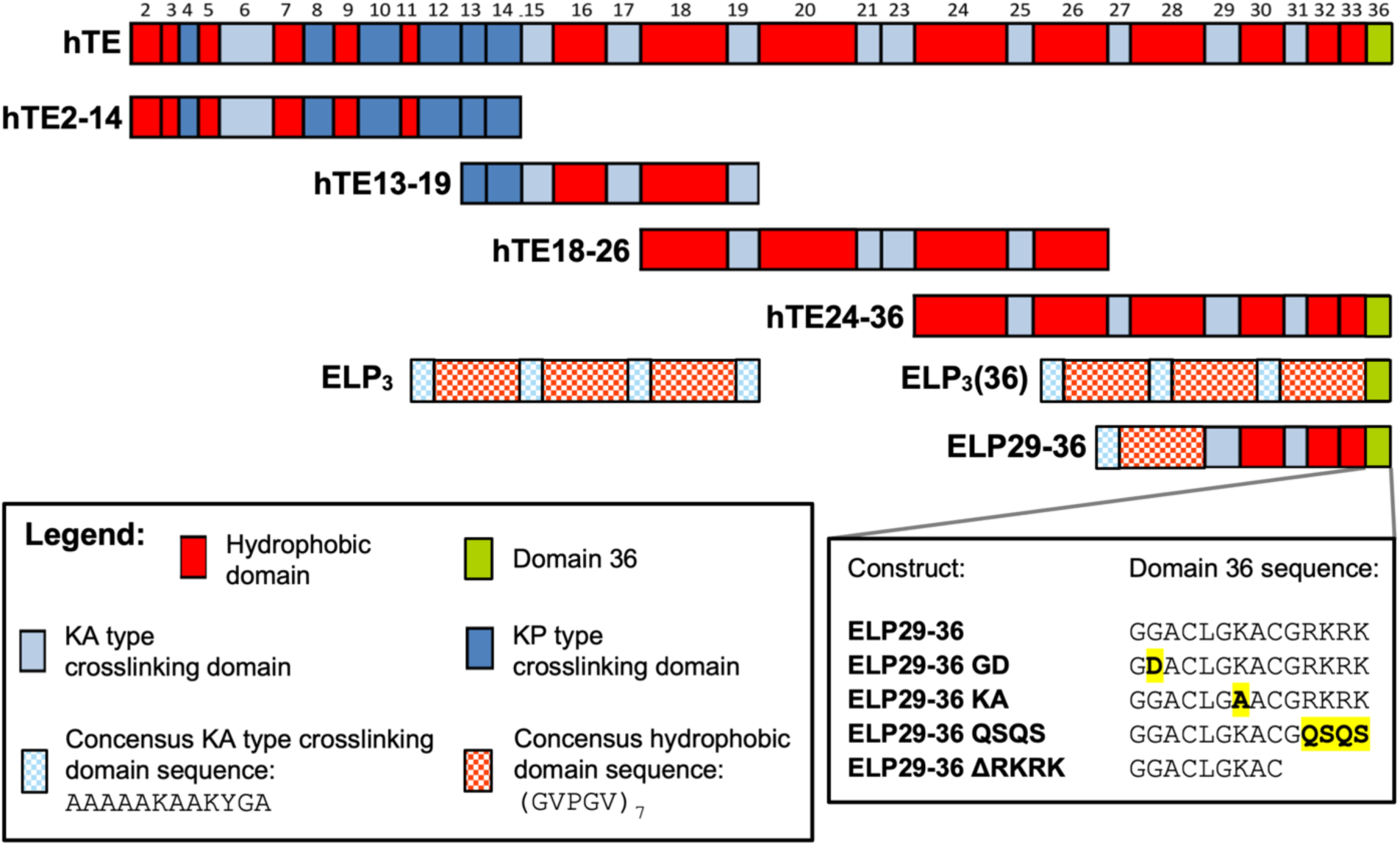
Schematic of tropoelastin domain organization and ELP constructs used in this study. Human tropoelastin (hTE) is a ∼60 kDa protein with alternating hydrophobic (red) and crosslinking domain (blue) organization. The domain composition of shorter hTE fragments and ELPs are shown below the parent protein. Domain 36 (green) sequence and mutant domain 36 sequences are shown at bottom right. See SI Table S1 for detailed construct sequence parameters; see SI Table S2 for full construct sequences.

Domain 36 mutants were used to further investigate the role of domain 36 in the heparin interaction. The terminal charge cluster RKRK was deleted for hTE, ELP_3_(36), and ELP29-36 to produce the ΔRKRK mutants. Further domain 36 mutants were created within the ELP29-36 construct (Figure 1). The GD mutant (G685D as numbered within full tropoelastin) reduces flexibility and introduces a negative charge near the start of domain 36, and was suggested to confer structural and functional effects relevant to the pathogenesis of chronic obstructive pulmonary disease (COPD) [86].

The KA mutant is a mutation of the non-terminal lysine residue within domain 36, present between the two cysteine residues. The QSQS mutant replaces the positive RKRK with polar residues.

### Heparin promotes tropoelastin coacervation through charge interactions

To investigate the effects of heparin on ELP and tropoelastin phase separation, we measured coacervation temperatures (T_c_) for tropoelastin and ELPs with or without heparin at a range of NaCl concentrations. Samples were heated in a spectrophotometer cell and upon coacervation, droplet formation resulted in an increased absorbance at 440 nm. These data were used to obtain T_c_ as shown in Figure 2A. As previously reported for both tropoelastin and ELPs [20,87,88], the T_c_ decreases as a function of increased NaCl (Figure 2B). Upon addition of heparin, coacervation temperatures were reduced in a NaCl dependent manner, with heparin having no effect at higher NaCl concentrations (Figure 2B). These data are consistent with heparin-protein interactions being screened around 0.5–1 M NaCl [89–94]. Additionally, chemically desulfated heparin was less effective at promoting tropoelastin coacervation, resulting in smaller changes in T_c_ (Figure 2C). These data suggest that the primary effect of heparin on tropoelastin coacervation requires a charge-charge interaction driven by the GAG sulfate groups – on the cell surface, these interactions likely involve the sulfated domains of heparan [79]. The weak effect seen for desulfated heparin is likely due to the iduronic acid groups of heparin, which maintain a negative net charge on the GAG.

**Figure 2.**
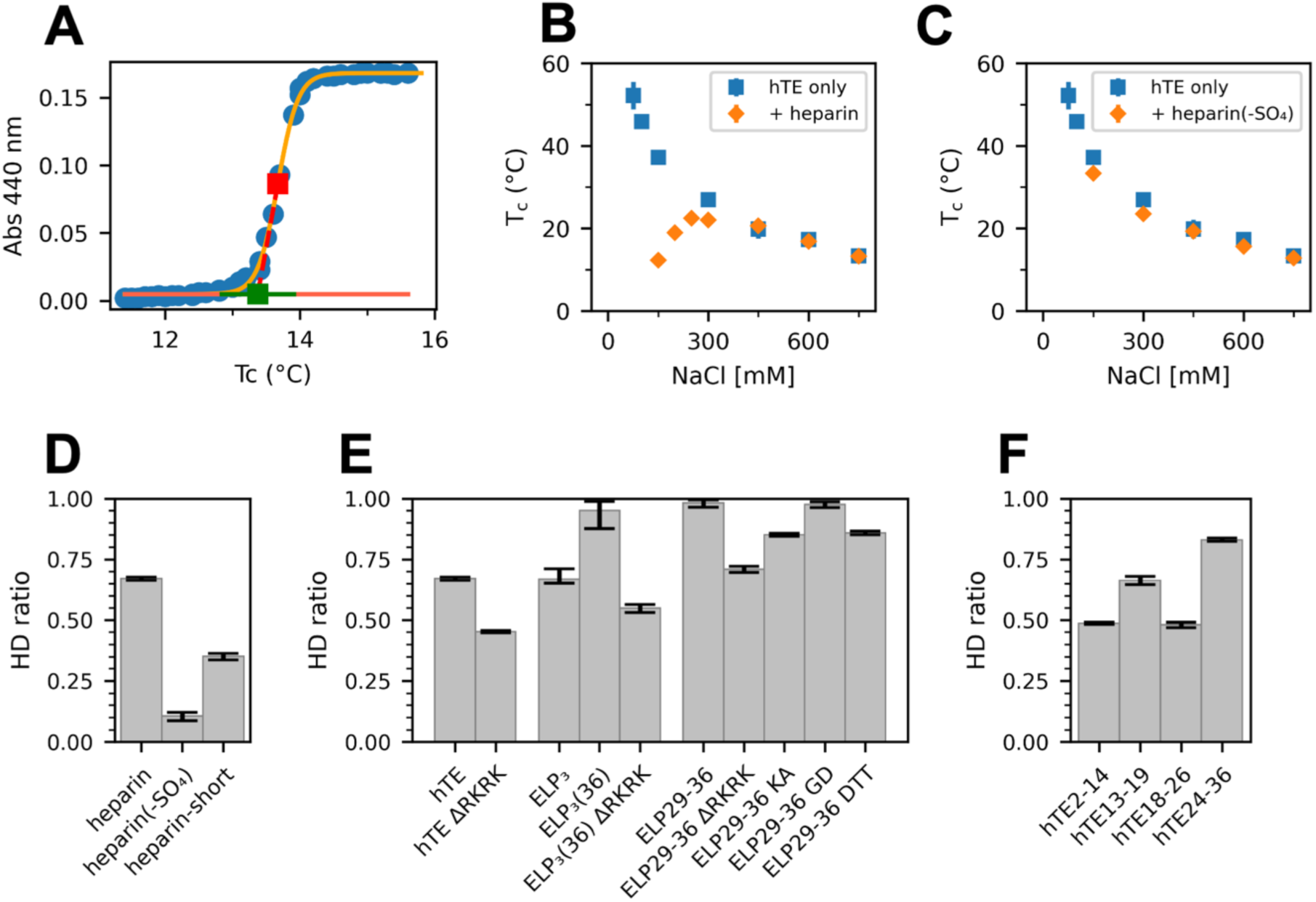
Heparin promotes tropoelastin coacervation through charged interactions. (A) Example tropoelastin turbidity measurement fit to a logistic curve (yellow, Eq. 1). The intersection of the tangent to the coacervation midpoint (red) and the pre-coacervation baseline (horizontal orange line) is taken as the coacervation initiation temperature (Tc, green dot and error bars). Coacervation temperatures measured as a function of NaCl concentration, in the presence and absence of GAG are shown here for (B) 25 µM hTE ± 6.25 µM heparin and (C) 25 µM hTE ± 6.25 µM heparin(-SO4) (desulfated heparin). The HD ratio (fractional change in Tc ± heparin at 150 mM NaCl, Eq. 3) for (D) tropoelastin with heparin, desulfated heparin, and a short fully sulfated heparin, (E) tropoelastin and ELPs containing variations in the domain 36 sequence, and (F) tropoelastin fragment constructs.

The strength of the heparin effect on T_c_ was quantified by calculating a heparin difference (HD) ratio as the fractional reduction in T_c_ due to the addition of heparin (Eq. 3). Values obtained at 150 mM NaCl were used to allow comparison of HD across all protein-heparin combinations (Figure 2D, SI Figures S1-S3). A lower HD ratio (reduced promotion of coacervation) for desulfated heparin supports a primarily charged interaction between GAG and protein. Reducing the size of a fully sulfated heparin from 18 kDa to 6.3 kDa also results in a lower HD ratio (Figure 2D, SI Figure S1A), suggesting that GAG binding to multiple tropoelastin molecules may play a role in promoting coacervation.

Sequence mutants support a role for domain 36 in binding heparin and promoting coacervation. The effect of heparin on tropoelastin lacking the C-terminal charge cluster in domain 36 (hTE ΔRKRK) is weaker than for the intact protein (Figure 2E, SI Figure S1B), suggesting an important role for the terminal charge cluster in domain 36. The effect of domain 36 was further confirmed using ELPs. ELP_3_ has a lower HD ratio than ELP_3_(36) (Figure 2E), and the ΔRKRK mutants of both ELP29-36 and ELP_3_(36) result in a lower HD ratio and weaker heparin effect (Figure 2E), supporting a role for the positive charge cluster in domain 36–heparin interaction. Removing a single positive charge (KA mutant of ELP29-36) also reduces the HD ratio, although adding a negative charge (GD mutation in domain 36) does not (Figure 2E). Reduction of the disulfide bond in domain 36 also reduces the heparin effect for ELP29-36 (Figure 2E), suggesting that the structure of domain 36 may also play a role in the interaction with heparin.

Importantly, there is not a linear relationship of the HD ratio with the tropoelastin or ELP construct net charge per residue (SI Figure S4), suggesting that total charge is not the only sequence feature important for interaction with heparin. In particular, there are constructs which have the same or similar charge densities but have different HD ratios, such as the ELP29-36 KA and GD mutants, ELP29-36 without or with reduced domain 36, and hTE and hTE2-14. Comparing the tropoelastin fragment constructs, hTE2-14 has a lower than expected HD ratio given its charge density, which may again reflect a role for local structural properties in heparin binding – KP crosslinking domains, as predominantly found in domains 2-14, are proline rich and thus unable to adopt the transient α-helical structures adopted by alanine-rich KA domains. Meanwhile, hTE13-19 has a higher than expected HD ratio, consistent with previous studies identifying heparin binding regions around crosslinking domains 15 and 17 [75,76].

### Heparin acts like a nucleating agent in promoting ELP coacervation

Our data so far strongly supports a primarily charge-based interaction between tropoelastin and heparin, so we hypothesized that there may be an optimal binding stoichiometry based on charge neutralization. We investigated this hypothesis by measuring coacervation of ELP29-36 as a function of heparin concentration (Figure 3). In addition to the coacervation initiation temperature T_c_, we also observe changes in the cooperativity of coacervation (approximated as the rate of absorbance increase which is reported as the logistic growth rate parameter, *k*; Eq. 1). This is a somewhat qualitative measure since absorbance during coacervation is affected by both droplet size and number, which may be solution condition dependent [95,96].

**Figure 3.**
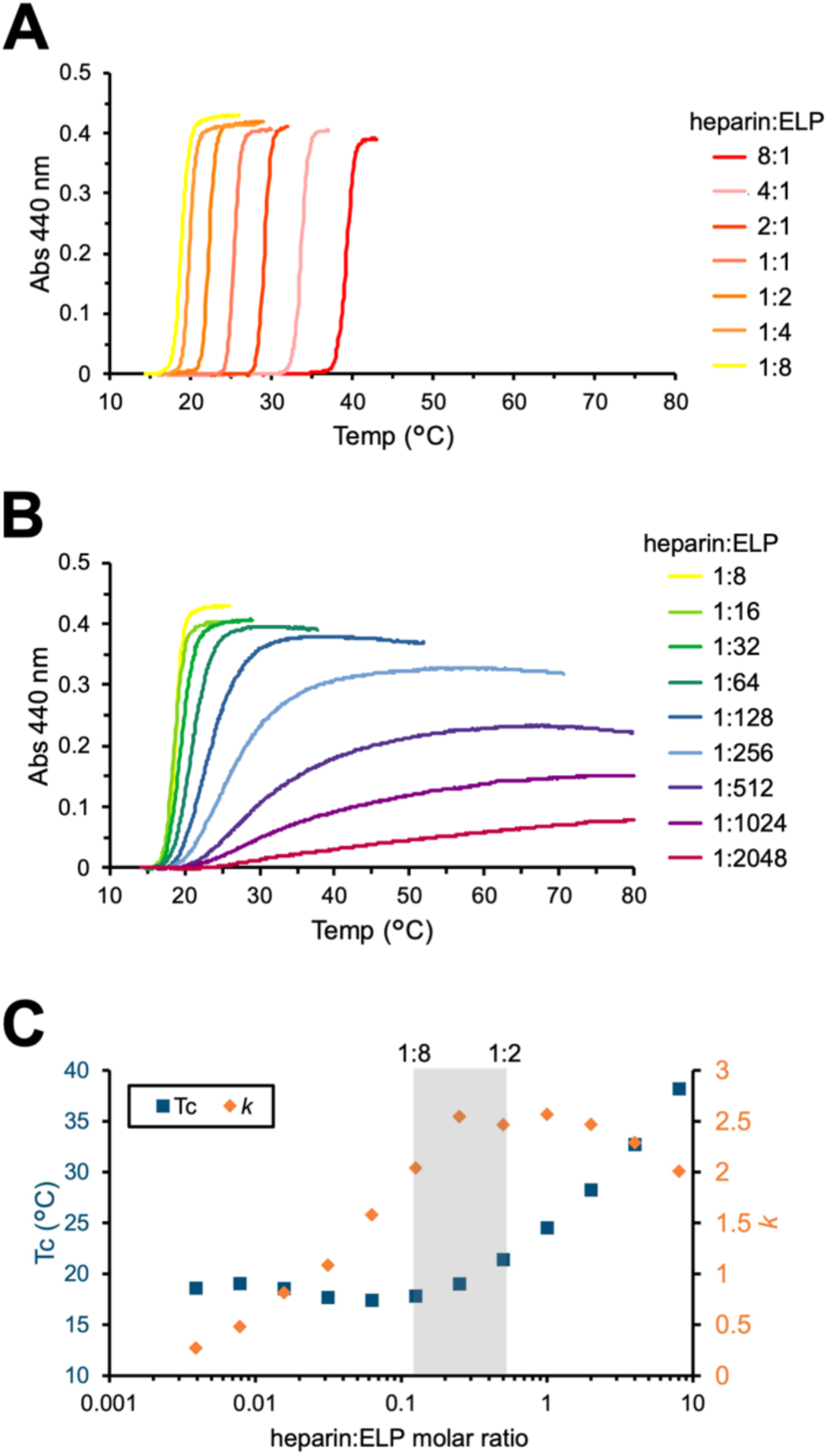
The ratio of heparin:protein changes both the initiation temperature and rate of ELP29-36 coacervation. Coacervation of ELP29-36 (200 µM) was triggered at 150 mM NaCl with varying concentrations of heparin, shown here as the heparin:ELP molar ratio. For visibility, data were separated into two sets, with (A) containing data for heparin:protein ratios of 8:1 to 1:8 and (B) showing ratios from 1:8 to 1:2048. (C) Comparison of Tc and k values obtained from fitting the coacervation curves for each heparin:ELP ratio. Note that data for samples with heparin:ELP ratios of 1:512, 1:1024, and 1:2048 were omitted from the final analysis due to poor fits with Eq. 1.

At heparin:ELP ratios higher than 1:8 there is an increase in T_c_ (Figure 3A) while at lower ratios T_c_ is relatively constant but there is a slowed progress of the coacervation event as the amount of heparin present is reduced (Figure 3B). The combined analysis of initiation (T_c_) and rate (*k*) of coacervation shown in Figure 3C highlights these trends and allows identification of conditions ideal for rapid favorable coacervation to occur. The heparin used here has a molecular weight of 18 kDa, which with an average of 3 sulfate groups per disaccharide subunit (each subunit is ∼660 Da, for ∼27 total disaccharides per chain) would give a net charge of approximately -104. ELP29-36 has a net charge of +12, so charge neutralization would occur when each heparin interacts with 8–9 molecules of ELP. This approximation closely matches the coacervation data shown in Figure 3, with increased heparin competing for ELP binding, resulting in higher T_c_ and lower *k*. This suggests that heparin is acting as a nucleating agent, bringing multiple ELPs together to promote coacervation. At reduced heparin (here, less than 1:16), the slower rate of coacervation likely reflects either a reduced number of initiating nuclei in the sample, a limited rate of coacervate droplet growth, or both.

It is important to note that the increase in coacervation temperatures with higher heparin:ELP ratios is not an inhibition of coacervation but rather a weakening of the heparin effect. ELP29-36 coacervation without heparin at 150 mM NaCl is expected to occur at extremely high temperatures or be impossible due to charge repulsion. Heparin significantly reduces the coacervation temperature, and high heparin:ELP ratios leads to a slightly smaller reduction.

### Direct observation of heparin binding to tropoelastin domain 36 and KA crosslinking domains

The site-specific details of heparin binding to ELP_3_, ELP_3_(36), hTE13-19, and ELP29-36 were investigated using solution NMR. All experiments were performed under non-coacervating conditions at 10 °C. Backbone ^1^H-^15^N HSQC spectra of all constructs show a reduced dispersion of ^1^H chemical shifts (7.9–8.8 ppm, Figure 4, SI Figure S5), indicative of intrinsic disorder and are consistent with previous NMR studies of elastin-like sequences [19,97,98]. ELP_3_ and ELP_3_(36) amide ^1^H and ^15^N chemical shifts were assigned using a strategy we have previously reported for ELP_3_ [19]. For these constructs, the repetitive sequence elements give rise to a single set of peaks, reflecting the disordered and polymer-like nature of these segments. hTE13-19 and ELP29-36 backbone chemical shifts were assigned using standard 3D NMR experiments as described elsewhere [82], and amide ^1^H and ^15^N chemical shifts with heparin were assigned to approximately 50-60%.

**Figure 4.**
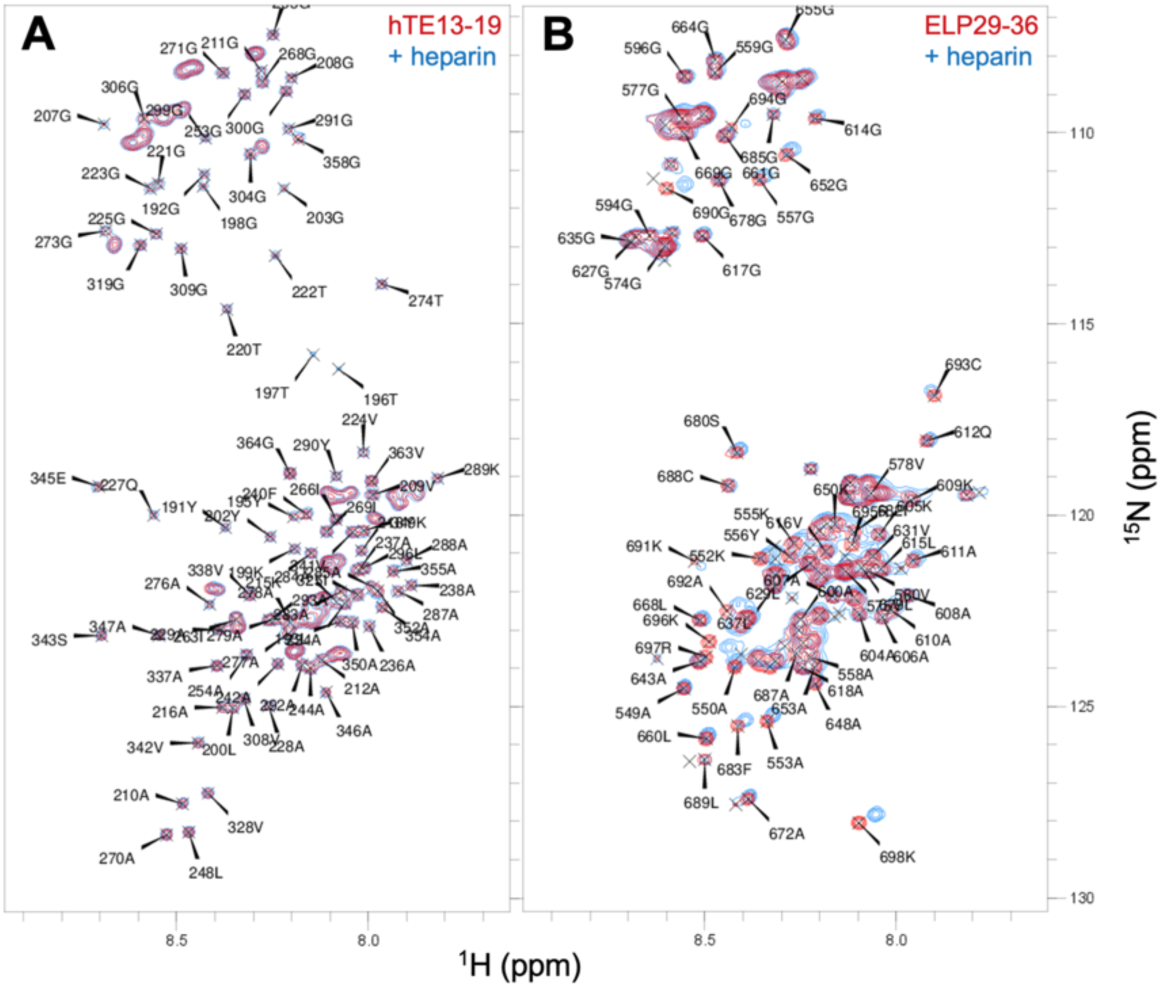
Backbone chemical shift changes upon addition of heparin to hTE13-19 or ELP29-36. ^1^H-^15^N HSQC spectra of (A) hTE13-19 and (B) ELP29-36 at 200 µM without (red) or with heparin at 50 µM (blue). The labels are the assigned residues, numbered with domain 36 aligned to its position within full tropoelastin with the last residue being Lys 698. Buffers contain 100 mM NaCl, 50 mM sodium phosphate, pH 7.2. All spectra were recorded at 700 MHz at 10 °C under non-coacervating conditions.

Upon addition of heparin, a subset of ^1^H-^15^N resonances exhibited a heparin-dependent change in chemical shift (Figure 4, SI Figures S5–S7). The observation of a single sharp resonance at all heparin concentrations suggests that there is fast exchange between the bound and unbound states. In all cases, there is little change in secondary structure propensity upon heparin binding, as predicted from chemical shift data using δ2D [99]. This is shown for ELP29-36 in SI Figure S8, with 85% of assigned residues exhibiting less than 5% total change in propensity using δ2D [99] and no indication of correlated local structural change. Overall, the largest changes in chemical shift are observed for KA crosslinking domains and domain 36, which contain pairs of lysine residues and the terminal RKRK motif, respectively (Figure 5, SI Figure S7).

**Figure 5.**
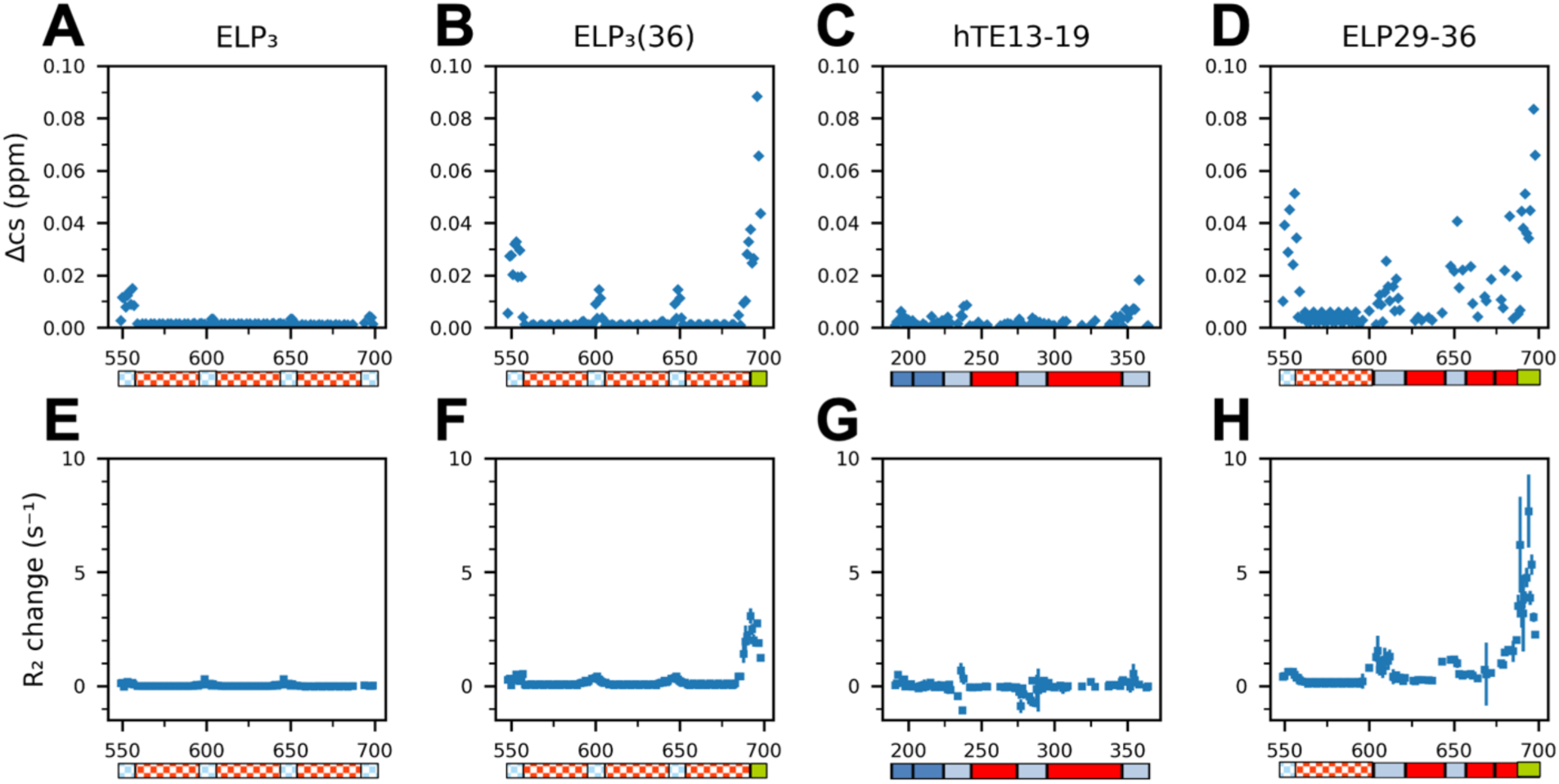
Changes in amide chemical shifts and 15N R2 for ELPs in the presence of heparin. ^1^H-^15^N amide backbone chemical shift change (Δcs) upon addition of 50 µM heparin to 200 µM of (A) ELP_3_, (B) ELP_3_(36), (C) hTE13-19, and (D) ELP29-36. A schematic of each construct’s domain organization is aligned below each plot, coloured to match Figure 1. ^15^N R_2_ relaxation rate change upon heparin addition is shown for (A) ELP_3_, (B) ELP_3_(36), (C) hTE13-19, and (D) ELP29-36. ^15^N R_2_ relaxation rates were acquired at 600 MHz for ELP_3_ and hTE13-19 and at 700 MHz for ELP_3_(36) and ELP29-36. All experiments were performed with buffer containing 100 mM NaCl, 50 mM sodium phosphate pH 7.2, 10% D2O, at 10 °C under non-coacervating conditions.

To further investigate the binding of heparin to ELPs, ^15^N amide R_1_ and R_2_ relaxation rates were used to monitor changes in local and global backbone dynamics (SI Figure S9). In the presence or absence of heparin, R_1_ rates remained relatively uniform along all protein chains, suggesting limited changes in fast timescale motion (ns–ps) upon GAG binding. For all constructs, amide R_2_ values were higher for crosslinking domains relative to the hydrophobic domains, likely due to the presence of transient helical structures in KA type crosslinking domains. This is more pronounced in both hTE13-19 and ELP29-36, suggesting increased helical stability in these constructs. Upon addition of heparin, R_2_ relaxation rate is most increased for residues in domain 36 and in the crosslinking domains of the domain 36 containing proteins (Figure 5). In contrast, very small changes in R_2_ rates are observed for ELP_3_ and hTE13-19. These data suggest that heparin binding is strongest in domain 36, the presence of which can promote or stabilize further interactions with nearby crosslinking domains.

To confirm the transient nature of the heparin-protein interactions, the hydrodynamic radius (R_h_) of each ELP was measured in the presence and absence of heparin, using NMR diffusion. Qualitatively the diffusion measurements show that increasing heparin concentration reduces the diffusion rate of all ELPs, with the greatest effect for domain 36 containing constructs ELP_3_(36) and ELP29-36 (SI Figure S10). Calculation of the R_h_ of ELPs shows a similar trend, with ELP29-36 exhibiting the largest R_h_ change from ∼3 to 4 nm (SI Figure S10). The reported R_h_ for heparin ranges from 2.5–4 nm [100–102], such that a stable ELP-heparin complex would have an expected R_h_ between 4 and 7 nm. Our chemical shift data suggest fast exchange between the bound and unbound states and the measured R_h_ values in the presence of heparin are smaller than expected for a stable complex, supporting a transient interaction. While we do not observe a significant reduction in NMR signal intensity, we cannot definitively exclude the presence of a minor stable complex in the heparin containing samples.

### Heparin brings ELPs into closer proximity pre-coacervation

The data presented above suggest the formation of transient heparin-ELP complexes that act to lower the energy barrier for coacervation. Such a nucleation or scaffolding behaviour is expected to bring multiple ELPs together on a single heparin chain. Paramagnetic relaxation enhancement (PRE) NMR was used to probe for new intermolecular and intramolecular protein-protein interactions formed upon addition of heparin to ELP29-36 (Figure 6, SI Figure S11). In PRE experiments, proximity to a paramagnetic label (here MTSL) will reduce amide peak intensity in a distance-dependent manner, monitored as the changes between ^1^H-^15^N HSQC spectra for samples containing paramagnetic (oxidized) and diamagnetic (reduced) MTSL (I_ox_/I_red_ ratio).

**Figure 6.**
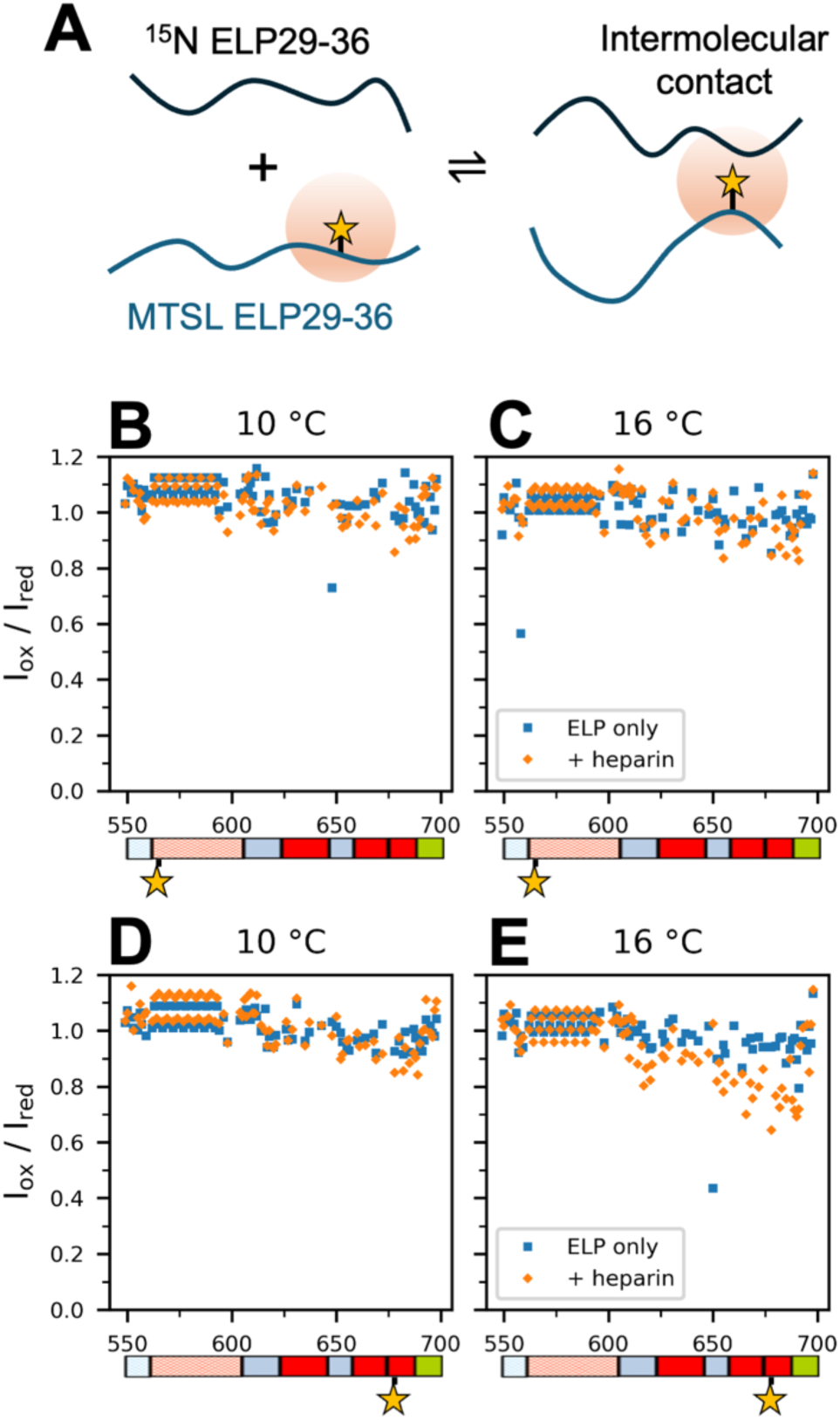
**PRE NMR data obtained for MTSL-labelled ELP29-36 as a function of heparin and sample temperature. (**A) Schematic of a sample to observe intermolecular PRE contacts, a 50:50 mixture of ^15^N ELP29-36 with unlabelled, MTSL tagged ELP29-36 (100 µM each, 200 µM total). [Top row] I_ox_/I_red_ data for ^15^N-ELP29-36 in the presence of N-terminally MTSL-labeled ELP29-36, ± heparin and at (B) 10 °C or (C) 16 °C. [Bottom row] I_ox_/I_red_ data for ^15^N-ELP29-36 in the presence of C-terminally MTSL-labeled ELP29-36, ± heparin and at (D) 10 °C or (E) 16 °C. For samples with heparin, the expected coacervation temperature is ∼18 °C. The domain organization of ELP29-36 and location of the MTSL tag is shown below each panel.

Samples containing a mixture of MTSL-tagged ELP29-36 and ^15^N labelled ELP29-36 were used to detect intermolecular interactions, while intramolecular interactions were probed using samples containing ^15^N-labelled, MTSL-tagged ELP29- 36. No significant change in I_ox_/I_red_ values were observed in the double-labeled samples (SI Figure S11), suggesting that heparin binding doesn’t promote intramolecular ELP-ELP contacts. PRE data obtained for the mixed ^15^N-labelled and C-terminally MTSL-tagged ELP29-36 samples show a reduction in peak intensity for residues near the C-terminus, but only at temperatures approaching T_c_ (Figure 6E). At lower temperatures, or with N-terminally MSTL-labelled protein, no change in I_ox_/I_red_ is observed. These data suggest that the transient domain 36–heparin interaction may be longer lived at higher temperatures and demonstrates the ability of heparin to bring multiple ELP chains together pre-coacervation.

## Discussion

The potential for glycosaminoglycans such as heparin to promote phase separation of tropoelastin is well established [40–42,70–72], however the molecular basis for this effect has not been previously elucidated. While specific interactions between heparin and either tropoelastin domain 36 or domains 15 and 17 have been proposed, [37,75,76] other studies have instead suggested a role for non-specific GAG-tropoelastin interactions, likely driven by charge interactions [42,71,77]. Our data provide evidence for a more complex mode of interaction in which non-specific interactions with multiple lysine containing CLDs, including 15 and 17, are present and sufficient for heparin-induced coacervation but with a much stronger effect in the presence of domain 36 (Figure 7). Specific binding of heparin to domain 36 appears to be driven by the highly conserved RKRK charge cluster [15] and results in a stronger reduction of coacervation temperature than would be expected based solely on the added local charge. Once brought into proximity of heparin by domain 36 interactions, CLD-heparin interactions along the chain become more favorable, resulting in transient multivalent interactions between GAG and protein. This is reminiscent of the fuzzy binding mode of integrin alphaVbeta3 with multiple domains of tropoelastin, including domain 36 [103]. Similar fuzzy complexes have also been proposed for tropoelastin interactions with MAGP [65,66] and fibulin 5 [104].

**Figure 7.**
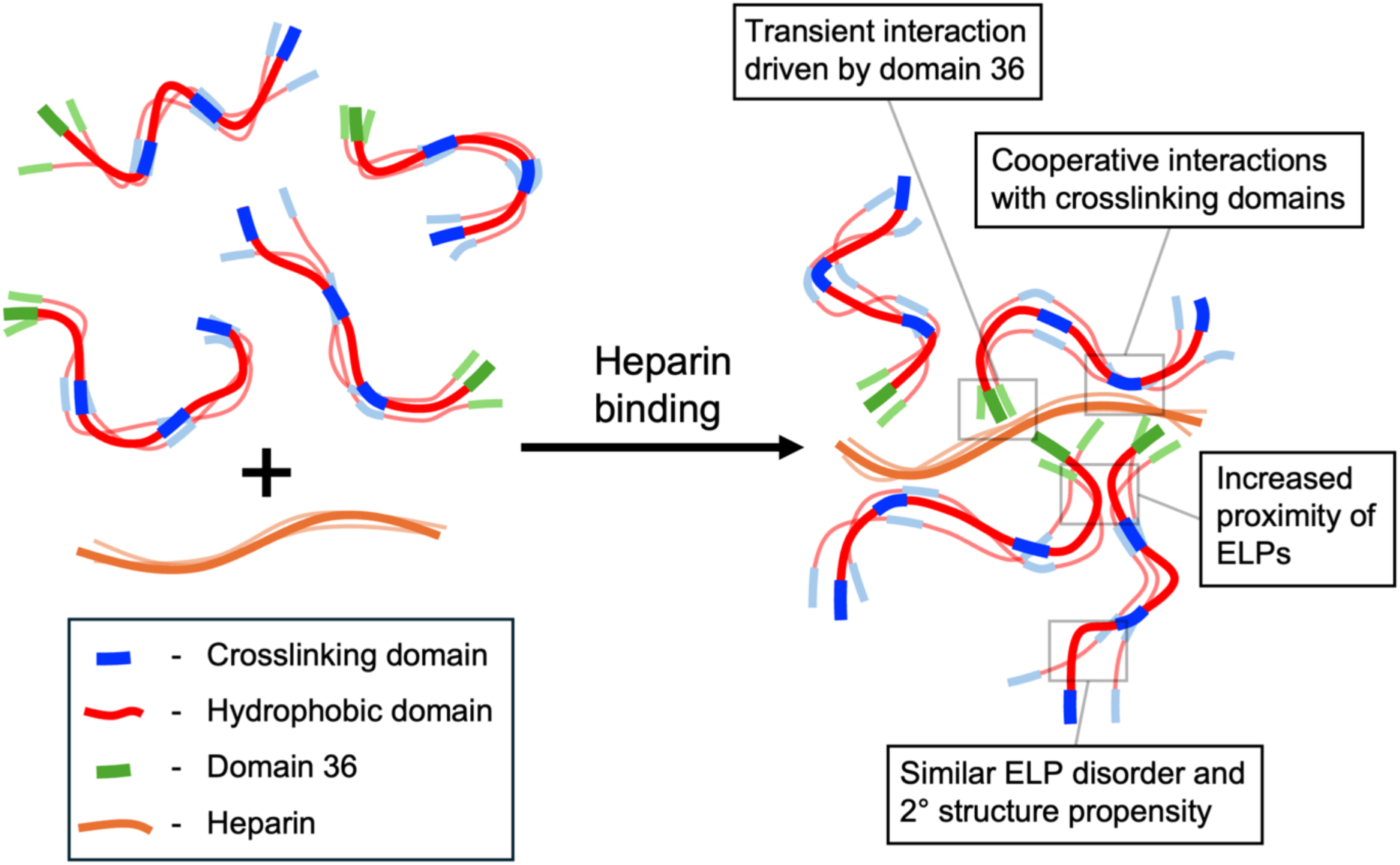
Model for the interaction of heparin and domain 36-containing ELPs. The interaction with heparin is transient and driven by tropoelastin domain 36 and the sulfate groups of heparin. Binding to domain 36 then results in cooperative weak interactions with crosslinking domains. Multiple ELPs bind to each heparin chain, creating a nucleating effect that reduces the energetic barrier to coacervation without altering the structural properties of the ELPs.

Since our NMR data suggest that interaction with heparin results in minimal changes in local secondary structure propensity, it is likely that heparin promotes phase separation simply through acting as a nucleating agent, bringing multiple tropoelastin/ELP molecules into close proximity, leading to the formation of transient clusters and reducing the energetic barrier to coacervation (Figure 7). These data are consistent with previous observations of transient ELP self-association pre-coacervation [105,106] and studies suggesting that mechanistically, the formation of pre-coacervation clusters can promote coacervation by increasing the nucleation rate and kinetics of coacervation [107–110]. While previous circular dichroism data indicated an increased α-helix propensity upon GAG binding to tropoelastin at physiological temperatures [41,78], our NMR data were obtained at 10 °C, where CLD helical structure is partly stabilized [111]. This suggests that while heparin binding may favour helical segments, induction of such is not likely a trigger for coacervation. Note that although we cannot exclude the possibility of more global shifts in the tropoelastin conformational ensemble, the similarity in coacervation behaviour of ELPs and the full-length protein suggest a similar mechanism for heparin in both systems.

Macroscopically, our data also suggest that different heparin:ELP ratios may influence the rate of growth of the droplet phase (ripening/coarsening) but are unclear on the mechanism, whether droplet growth may be diffusion or conversion-limited [28,112,113]. However, the existence of an optimal heparin:ELP ratio for promoting coacervation is consistent with previous studies of protein-RNA complex coacervates, where turbidity measurements show maximum coacervation when the mixture of the components is charge-neutral [114–116]. Overall, the relative strengths of homo and heterotypic interactions support a scaffold-client relationship [31,117,118], where tropoelastin is acting as a scaffold and heparin is acting as a client to promote coacervation at low NaCl concentration.

Supporting its role in promoting coacervation, domain 36 is known to be required for proper elastic fibre assembly in humans, with mutations in this region resulting in impaired elastic fibre assembly [38,119], leading to congenital diseases such as autosomal-dominant cutis laxa [2,120–122] and increased susceptibility to late-onset diseases such as chronic obstructive pulmonary disease [86]. Domain 36 has previously been implicated as a key interaction site between tropoelastin and other ECM components, including integrins [123,124], MAGP [38,64], and fibrillin [38,58]. Interestingly, domain 36 is modified or removed in the mature elastic fibre, suggesting that its role is limited to earlier steps in the assembly process, rather than stabilizing the mature fibre or modulating elasticity [86,125]. This is in keeping with our data showing that domain 36 is a key driver of GAG-tropoelastin interactions that promote assembly.

Overall, our results provide evidence for a molecular mechanism through which extracellular GAGs can promote tropoelastin coacervation at physiological ionic strength, supporting a role for GAG-tropoelastin interactions at the cell-surface in initiating elastic fibre assembly. GAG length, sulfation, and concentration influence their ability to promote tropoelastin coacervation, suggesting that the type and amount of GAG present in different tissues are likely major factors involved in modulating tropoelastin coacervation and elastic fibre assembly. Given the distribution of different GAG types in the ECM [1,126] and previous studies showing different types of GAGs promoting tropoelastin coacervation [40–42], interactions with many ECM GAGs likely influence elastic fibre assembly as tropoelastin coacervation is initiated on the cell surface and coacervates are transported through the ECM to the microfibril. GAGs interact differently with different regions of tropoelastin, potentially influencing tropoelastin crosslink type and distribution, thus affecting elastic properties of the mature fibre [35,127,128]. The gradual crosslinking of tropoelastin, which eliminates the positive charges within crosslinking domains, is likely to reduce GAG interactions, suggesting that non-charged interactions with other ECM components such as fibulins or fibrillin may be more important in controlling the later stages of elastic fibre assembly.

## Experimental Procedures

### Protein expression and purification

All ELPs and full-length human tropoelastin were expressed in BL21 (DE3) *E. coli* as described previously [19,29,111]. Transformed cells were grown in 1 L of Terrific Broth containing 100 μg/mL ampicillin and 0.8 % glucose. Cells were grown shaking at 37 °C until a cell OD_600_ ≈ 1.0–1.1 and were induced with 200 μM IPTG. After 3.5 hours of protein expression, cells were pelleted by centrifugation. Isotopic labelling for NMR was achieved by growth in M9 minimal medium supplemented with ^15^N NH_4_Cl as the only nitrogen source or ^13^C glucose as the only carbon source.

For purification, cell pellets were dissolved in 20 mL of 80% formic acid with > 1 g cyanogen bromide (CNBr) incubated at room temperature overnight for ∼16 h. The CNBr solution was then dialysed with a 3.5 kDa cutoff into 20 mM sodium acetate pH 5.25, after which the cell debris was pelleted by centrifugation at 12000 x g for 15 min. The filtered supernatant was then applied to cation exchange FPLC, with the polypeptides of interest eluting around 15–20 mS/cm; constructs with domain 36 eluted around 25–30 mS/cm. The selected FPLC fractions were then dialysed into water and lyophilized before a final step of C4 reverse phase HPLC, then lyophilized again to obtain the final, dry polypeptide.

### Turbidometry

Coacervation of ELPs and tropoelastin was measured in a cuvette format using a Shimadzu UV-2401PC UV-vis spectrometer. To prepare samples, dry ELP/tropoelastin was dissolved in 200 mM tris pH 7.5 on ice, and heparin was dissolved in water at room temperature for at least 30 minutes. Then, samples were centrifuged at 20,000 rcf to remove aggregates. The supernatant was then mixed on ice with concentrated NaCl stock solutions to produce the final samples, which contained 50 mM tris pH 7.5 and varying concentrations of NaCl. For coacervation experiments, samples were heated at a rate of 2 °C/min, starting at 12 °C and ending approximately 10 °C past the initiation of coacervation, up to 95 °C. Coacervation was monitored using absorbance at 440 nm. Protein concentrations were 200 μM for ELPs and shorter tropoelastin fragment constructs, and 25 μM for full-length tropoelastin and tropoelastin ΔRKRK. Heparin for coacervation experiments was added to give a 1:4 heparin:protein molar ratio (i.e. 50 µM for ELPs and tropoelastin fragment constructs, 6.25 µM for tropoelastin). For hTE coacervation with hep-short, the molarity was increased by 2.85 times (i.e. 17.8 µM short heparin) to maintain similar levels of net charge from heparin, thus testing only the effect of changing heparin length.

Fitting of the coacervation initiation temperature was done using custom Python code using a standard logistic function (Eq.1).

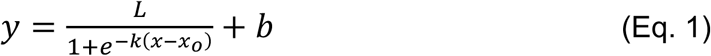

L is the amplitude or height, *k* is the logistic growth rate, *x_0_* is the midpoint (sometimes referred to as the Tm), *b* is the vertical shift, *x* is the temperature and *y* is the absorbance. The data from the spectrometer was first pre-processed by removing data past the peak of coacervation and applying a linear correction to flatten a sloped baseline if necessary. Fitting was performed using curve_fit from scipy.optimize. The coacervation initiation temperature (T_c_) is the x-coordinate of the intersection of the lower baseline and the tangent line from the midpoint of the logistic growth curve. Reported coacervation temperatures are the average of three replicates unless specifically noted. The reported error is a quadrature sum of the sample standard deviation of the fitted temperatures and the propagated fit error.

The heparin used throughout this study (18 kDa) was from Sigma-Aldrich (from porcine intestinal mucosa). Desulfated heparin was produced using a protocol from Seto et al. [129]. Briefly, heparin (18 kDa from Sigma-Aldrich) was dissolved at 11.1 mg in 2.2 mL of methanol with 0.5% v/v acetyl chloride and stirred for 7 days. On days 1, 4, and 6, the heparin was centrifuged at ∼20,000 rcf for 15 minutes, then the pellet resuspended in new acidic methanol. On day 7, the methyl ester of heparin was centrifuged and resuspended in 200 µL of water, then split into two tubes, and washed with 2 mL ethanol each, centrifuging and resuspending 3 times. The final wash was with 2 mL ethyl ether, with the tubes combined, and then the heparin was freeze-dried to obtain the final product. Desulfation was verified using the dimethylmethylene blue (DMMB) assay [130]. Briefly, DMMB color reagent was prepared by mixing 16 mg DMMB (powder) in 1L distilled water with 3.04 g glycine and 2.37 g NaCl, stirred overnight wrapped in foil, then pH adjusted the next day to pH 3. Heparin and desulfated heparin standard samples were prepared by serial dilution, with final amount per well ranging from 0.00061–1.25 µg/well. DMMB color reagent was added, and absorbance read at 525 nm in a Synergy Neo2 plate reader (SBC Facility, The Hospital for Sick Children). Short heparin (6.3 kDa) was purchased from Iduron.

### Calculation of the heparin difference (HD) ratio

At low NaCl concentrations, many samples did not undergo coacervation in the absence of heparin within the experimental range of the spectrophotometer. To allow comparison of the T_c_ in the presence/absence of heparin, the T_c_ vs NaCl data were fit to allow extrapolation to lower NaCl concentrations and calculate the heparin difference (HD) ratio at 150 mM NaCl, near physiological salt concentration [131,132].

The T_c_ values obtained at a range of NaCl concentrations for a particular construct were fit using a modified Langmuir binding isotherm [88] (Eq. 2).

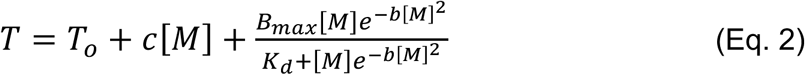

T_o_ is the coacervation temperature; c is a constant with units of temperature/molarity; B_max_ is a constant with units of temperature; K_d_ is a constant; b is a constant with units of inverse molarity. Fitting was done using the lmfit module in Python, iterating through five rounds using the default Levenberg-Marquardt (“leastsq”) method, with the result of each fit kept as the initial input for the subsequent round of fitting. To optimize the initial parameter values, a range of T_o_ and B_max_ values were tested while keeping the other parameters, c, K_d_, and b fixed, and selecting the T_o_/B_max_ values which produced the fit with the lowest reduced Χ^2^ (chi squared). The result of the optimized fit was then used as the initial guess for final fitting using the emcee method within lmfit, which samples the posterior distribution of the fit parameters using Markov Chain Monte Carlo, estimating a probability distribution for each parameter (see the online documentation for more details: https://lmfit.github.io/lmfit-py/fitting.html). Using the default settings, 100000 chains (parameter estimates) were produced (steps=1000, nwalkers=100). The chain (set of parameters) with the highest probability (the best fit) was used to extrapolate T_c_ to 150 mM NaCl. The distribution of parameters was used to estimate the error of the extrapolated point, selecting only the good fits, if the reduced Χ^2^ of the fit from the parameter values of the chain was within 0.05–0.95 critical value for the reduced Χ^2^ distribution with the given degrees of freedom. If no fits had reduced Χ^2^ between 0.05–0.95, then the top 50% of chains (50000 chains) with the highest probability were selected. The standard deviation of the selected fits was used as the error of the extrapolated T_c_.

The HD ratio was calculated as the ratio of the reduction in T_c_ caused by heparin and the protein only coacervation temperature at 150 mM NaCl (Eq. 3).

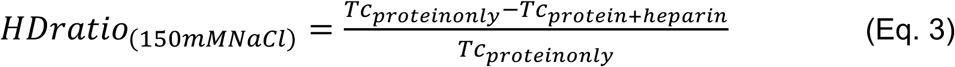

The HD ratio ranges between 0 (no effect of heparin on T_c_) and 1, with a reported error propagated from the standard deviation of measured T_c_. For ELPs where the protein only coacervation temperature was extrapolated to low NaCl concentration, the standard deviation of the fit error estimated from the emcee method was used in place of measurement error.

### Solution NMR

Solution NMR experiments were performed on Bruker Avance III spectrometers operating at a ^1^H frequency of either 600 or 700 MHz using a 5 mm TXI triple resonance probe. Standard Bruker experiments for chemical shift assignment were performed at 10 °C under non-coacervating conditions, including ^1^H-^15^N HSQC, HNCO, and HNcaCO. NMR sample buffers contained 50 mM sodium phosphate pH 7.2, 100 mM NaCl, 10% D_2_O. ELP_3_ and ELP_3_(36) amide ^1^H and ^15^N chemical shifts with and without heparin were assigned fully by copying overlapping resonances using previously assigned spectra from our lab [19]. hTE13-19 and ELP29-36 amide ^1^H and ^15^N resonances without heparin were fully assigned as described elsewhere [82]. From resolved peaks in the ^1^H-^15^N HSQC spectra, hTE13-19 amide ^1^H and ^15^N chemical shifts with heparin were assigned to 49% and ELP29-36 amide ^1^H and ^15^N chemical shifts with heparin were assigned to 57%. To predict secondary structure propensity with δ2D [99], for ELP29-36 200 µM with heparin 25 µM, HNCO and HNcaCO spectra were acquired in addition to the ^1^H-^15^N HSQC, assigning amide proton and nitrogen resonances to 76% and carbonyl carbon resonances to 84%. The total secondary structure propensity change is the sum of the change in α-helix, β-sheet, coil, and PPII propensity change. Chemical shift change was calculated using weighted differences in measured chemical shift, combined and weighted for each element as shown in Eq. 3, where Δ𝑐𝑠 is the total chemical shift change upon addition of heparin, and 𝛿_*H*_ and 𝛿_*N*_ are the chemical shift changes for the backbone amide ^1^H and ^15^N resonances. The suggested scaling factors α = 0.14 for ^15^N and α = 0.2 for ^15^N glycines were used [133].

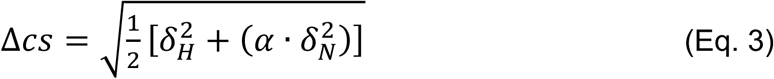

Heparin titrations were performed using 200 μM ELP with heparin concentrations ranging from 0–200 μM. PFG diffusion data was acquired using a stimulated echo experiment with bipolar gradients and WATERGATE solvent suppression (Bruker TopSpin sequence stebpgp1s19), with 0.2% dioxane added to the buffer as an internal control for sample viscosity. Sixteen ^1^H spectra were acquired at varying gradient strengths between 2% and 90% of the maximum output while keeping delay times constant. The diffusion coefficient was obtained by fitting peak intensity change to a standard equation for bipolar gradients, then converted to hydrodynamic radius (R_h_) by the Stokes-Einstein law [134]. The ^15^N relaxation rates R_1_ and R_2_ were determined using standard inversion recovery or spin-echo experiments (Bruker TopSpin sequences hsqct1etf3gpsi3d and hsqct2etf3gpsi3d); ten spectra were acquired with delay times between 50–1500 μs for R_1_ and 17–544 μs for R_2_ experiments. Least-squares fitting of signal attenuation was performed in ccpnmr using standard parameters.

To produce MTSL labelled samples for PRE experiments, a cysteine mutation was added as the MTSL attachment site, replacing a hydrophobic amino acid. For ELP29-36, the N-terminal mutation was 1V5C, replating the 5^th^ valine in the 1^st^ hydrophobic domain repeat. The C-terminal mutation was F677C. Domain 36 cysteines were also mutated to serine (C688S, C693S). Purified and dry protein was dissolved in a buffer containing 50 mM sodium phosphate pH 7, 50 mM NaCl, to a final concentration of 200–250 µM in a final volume of 1.7 mL. Dithiothreitol (DTT) was added to the protein to a final concentration of 2 mM. The protein + DTT mixture was incubated at room temperature for > 1 hr to fully reduce the cysteines. DTT was then removed using a desalting column using the manufacturer’s spin protocol (Cytiva PD-10 MiniTrap with sephadex G-25), eluting ∼1.7 mL per sample. After desalting, 35 µL of MTSL (from a 100 mM solution in acetonitrile) was added, to final MTSL concentration of 2 mM. Samples were wrapped in foil and incubated on a rocking stage overnight at room temperature. The next day, samples were dialyzed in water to remove excess MTSL, then lyophilized.

Samples for intermolecular PRE experiments were prepared using a 1:1 mixture of ^15^N labelled ELP and unlabelled ELP with MTSL tag (100 µM each, 200 µM total ELP concentration). Samples for intramolecular PRE experiments were prepared using a 1:1 mixture of ^15^N labelled ELP with MTSL tag and unlabelled ELP (100 µM each, 200 µM total ELP concentration). Samples with heparin contained 50 µM heparin (total heparin:ELP molar ratio 1:4). Buffer for PRE samples contained 50 mM sodium phosphate pH 7, 100 mM NaCl, and 10% D_2_O.

To perform the PRE experiments, first, an HSQC was acquired of sample with an oxidized (paramagnetic) MTSL label. Then, ascorbic acid was mixed into the sample to a final concentration of 2 mM and incubated for 2 hours to allow for complete reduction of MTSL before acquiring a second HSQC with reduced (diamagnetic) MTSL. Peaks were assigned, and peak intensities from the two HSQCs were used to produce I_ox_ and I_red_ and calculate the strength of the PRE effect I_ox_/I_red_. Spectra were acquired at 10 and 16 °C, using both N-terminal and C-terminal ELP29-36 MTSL tags, with and without heparin.

## Supporting Information

SI Tables S1 and S2, containing protein amino acid sequences and properties; SI Figures 1–11, containing additional coacervation and NMR data for ELP constructs.

## Acknowledgements

The authors acknowledge The Hospital for Sick Children’s Structural and Biophysical Core (SBC) Facility for providing experimental guidance and instrumentation.

## Funding Sources

This work was supported by a Grant in Aid from the Lung Health Foundation and a Grant in Aid from the Heart and Stroke Foundation of Canada Grant in Aid (grant number G-18-0022249).

## Author contributions

Robert Lu: Methodology, Investigation, Writing – Original Draft, Writing – Review & Editing; Sean Reichheld: Investigation, Resources, Writing – Review & Editing; Min Jin: Investigation, Writing – Review and Editing; Simon Sharpe: Conceptualization, Writing – Review & Editing, Supervision, Funding Acquisition.

## Notes

The authors declare no conflict of interest.

### Competing Interest Statement

The authors have declared no competing interest.

