## Supplemental Tables and Figures for "Heparin nucleates and promotes tropoelastin coacervation through a transient interaction driven by domain 36"

**Supplementary Information**

Robert Lu[a], Sean Reichheld[b], Min Jin[a], Simon Sharpe[a,b,*]

[a] Department of Biochemistry, University of Toronto, Toronto, ON, Canada, M5S 1A8

[b] Molecular Medicine Program, The Hospital for Sick Children, Toronto, ON, Canada M5G 0A4

### Table of Contents


***Supplementary Tables*3**

**SI Table 1. Sequence parameters of constructs used in this study.3**

**SI Table 2. Sequences of constructs used in this study.4**

***Supplementary Figures*5**

**SI Figure 1. Coacervation temperatures of hTE ± short heparin and hTE ΔRKRK ± heparin.5**

**SI Figure 2. Coacervation temperatures of ELP constructs with domain 36 mutants.6**

**SI Figure 3. Coacervation temperatures of tropoelastin fragment constructs.7**

**SI Figure 4. Plots of HD ratio against sequence net charge per residue7**

**SI Figure 5. Backbone $^{1}H$-$^{15}N$ HSQC upon addition of heparin for ELP3 and ELP3(36).8**

**SI Figure 6. Backbone $^{1}H$-$^{15}N$ HSQC upon titration of heparin for ELP3 and ELP3(36), zoomed in to the non-glycine region.9**

**SI Figure 7. Backbone $^{1}H$-$^{15}N$ chemical shift changes upon heparin titration.10**

**SI Figure 8. Predicted secondary structure propensity for ELP29-36 with heparin and change compared to no heparin.11**

**SI Figure 9. $^{1}H$-$^{15}N$ amide backbone R1 and R2 relaxation rates of ELPs ± heparin.12**

**SI Figure 10. Change in diffusion rate and estimated hydrodynamic radius of ELPs upon heparin titration.13**

**SI Figure 11. N- and C-terminal intramolecular PRE effect in ELP29-36 upon heparin binding and heating.14**

### Supplementary Tables

| Protein | Residues | kDa | neg_res | pos_res | net charge | cpm | NCPR | f- | f+ | FCR |
|---|---|---|---|---|---|---|---|---|---|---|
| hTE2-14 | 221 | 19.64 | 1 | 13 | 12 | 0.611 | 0.054 | 0.005 | 0.059 | 0.063 |
| hTE13-19 | 176 | 15.03 | 1 | 9 | 8 | 0.532 | 0.045 | 0.006 | 0.051 | 0.057 |
| hTE18-26 | 228 | 19.37 | 2 | 12 | 10 | 0.516 | 0.044 | 0.009 | 0.053 | 0.061 |
| hTE24-36 | 231 | 19.66 | 0 | 16 | 16 | 0.814 | 0.069 | 0.000 | 0.069 | 0.069 |
| hTE | 698 | 59.99 | 3 | 41 | 38 | 0.567 | 0.049 | 0.004 | 0.059 | 0.063 |
| hTE ΔRKRK | 693 | 59.36 | 3 | 37 | 34 | 0.573 | 0.049 | 0.004 | 0.053 | 0.058 |
| ELP29-36 | 152 | 13.09 | 0 | 12 | 12 | 0.917 | 0.079 | 0.000 | 0.079 | 0.079 |
| ELP29-36 ΔRKRK | 147 | 12.46 | 0 | 8 | 8 | 0.642 | 0.054 | 0.000 | 0.054 | 0.054 |
| ELP29-36 GD | 152 | 13.14 | 1 | 12 | 11 | 0.837 | 0.072 | 0.007 | 0.079 | 0.086 |
| ELP29-36 QSQS | 152 | 12.95 | 0 | 8 | 8 | 0.618 | 0.053 | 0.000 | 0.053 | 0.053 |
| ELP29-36 KA | 152 | 13.03 | 0 | 11 | 11 | 0.844 | 0.072 | 0.000 | 0.072 | 0.072 |
| $ELP_3$ | 153 | 12.80 | 0 | 8 | 8 | 0.625 | 0.052 | 0.000 | 0.052 | 0.052 |
| $ELP_3$(36) | 153 | 12.98 | 0 | 11 | 11 | 0.847 | 0.072 | 0.000 | 0.072 | 0.072 |
| $ELP_3$(36) ΔRKRK | 148 | 12.36 | 0 | 7 | 7 | 0.566 | 0.047 | 0.000 | 0.047 | 0.047 |

**SI Table S1. Sequence properties of protein constructs used in this study.** Residues – number of residues. kDa – molecular weight. neg_res – number of aspartic and glutamic acid residues. pos_res – number of lysine and arginine residues. net charge – overall net charge. cpm – charge per mass; net charge divided by molecular weight. NCPR – net charge per residue; net charge divided by number of residues. f+, f- – fraction of positively and negatively charged residues. FCR – fraction of charged residues.

| Protein | Sequence |
|---|---|
| hTE2-14 | GVPGAIPGGV PGGVFYPGAG LGALGGGALG PGGKPLKPVP GGLAGAGLGA GLGAFPAVTF PGALVPGGVA DAAAAYKAAK<br>AGAGLGGVPG VGGLGVSAGA VVPQPGAGVK PGKVPGVGLP GVYPGGVLPG ARFPGVGVLP GVPTGAGVKP KAPGVGGAFA<br>GIPGVGPFGG PQPGVPLGYP IKAPKLPGGY GLPYTTGKLP YGYGPGGVAG AAGKAGYPTG T |
| hTE13-19 | GGYGLPYTTG KLPYGYGPGG VAGAAGKAGY PTGTGVGPQA AAAAAAKAAA KFGAGAAGVL PGVGGAGVPG VPGAIPGIGG<br>IAGVGTPAAA AAAAAAAKAA KYGAAAGLVP GGPGFGPGVV GVPGAGVPGV GVPGAGIPVV PGAGIPGAAV PGVVSPEAAA<br>KAAAKAAKYG ARPGVG |
| hTE18-26 | AGVPGVGVPG AGIPVVPGAG IPGAAVPGVV SPEAAAKAAA KAAKYGARPG VGVGGIPTYG VGAGGFPGFG VGVGGIPGVA<br>GVPGVGGVPG VGGVPGVGIS PEAQAAAAAK AAKYGVGTPA AAAAKAAAKA AQFGLVPGVG VAPGVGVAPG VGVAPGVGLA<br>PGVGVAPGVG VAPGVGVAPG IGPGGVAAAA KSAAKVAAKA QLRAAAGLGA GIPGLGVGVG VPGLGVGA |
| hTE24-36 | GVGLAPGVGV APGVGVAPGV GVAPGIGPGG VAAAAKSAAK VAAKAQLRAA AGLGAGIPGL GVGVGVPGLG VGAGVPGLGV<br>GAGVPGFGAV PGALAAAKAA KYGAAVPGVL GGLGALGGVG IPGGVVGAGP AAAAAAAKAA AKAAQFGLVG AAGLGGLGVG<br>GLGVPGVGGL GGIPPAAAAK AAKYGAAGLG GVLGGAGQFP LGGVAARPGF GLSPIFPGGA CLGKACGRKR K |
| hTE | GGVPGAIPGG VPGGVFYPGA GLGALGGGAL GPGGKPLKPV PGGLAGAGLG AGLGAFPAVT FPGALVPGGV ADAAAAYKAA<br>KAGAGLGGVP GVGGLGVSAG AVVPQPGAGV KPGKVPGVGL PGVYPGGVLP GARFPGVGVL PGVPTGAGVK PKAPGVGGAF<br>AGIPGVGPFG GPQPGVPLGY PIKAPKLPGG YGLPYTTGKL PYGYGPGGVA GAAGKAGYPT GTGVGPQAAA AAAAKAAAKF<br>GAGAAGVLPG VGGAGVPGVP GAIPGIGGIA GVGTPAAAAA AAAAAKAAKY GAAAGLVPGG PGFGPGVVGV PGAGVPGVGV<br>PGAGIPVVPG AGIPGAAVPG VVSPEAAAKA AAKAAKYGAR PGVGVGGIPT YGVGAGGFPG FGVGVGGIPG VAGVPGVGGV<br>PGVGGVPGVG ISPEAQAAAA AKAAKYGVGT PAAAAAKAAA KAAQFGLVPG VGVAPGVGVA PGVGVAPGVG LAPGVGVAPG<br>VGVAPGVGVA PGIGPGGVAA AAKSAAKVAA KAQLRAAAGL GAGIPGLGVG VGVPGLGVGA GVPGLGVGAG VPGFGAVPGA<br>LAAAKAAKYG AAVPGVLGGL GALGGVGIPG GVVGAGPAAA AAAAKAAAKA AQFGLVGAAG LGGLGVGGLG VPGVGGLGGI<br>PPAAAAKAAK YGAAGLGGVL GGAGQFPLGG VAARPGFGLS PIFPGGACLG KACGRKRK |
| hTE<br>ΔRKRK | GGVPGAIPGG VPGGVFYPGA GLGALGGGAL GPGGKPLKPV PGGLAGAGLG AGLGAFPAVT FPGALVPGGV ADAAAAYKAA<br>KAGAGLGGVP GVGGLGVSAG AVVPQPGAGV KPGKVPGVGL PGVYPGGVLP GARFPGVGVL PGVPTGAGVK PKAPGVGGAF<br>AGIPGVGPFG GPQPGVPLGY PIKAPKLPGG YGLPYTTGKL PYGYGPGGVA GAAGKAGYPT GTGVGPQAAA AAAAKAAAKF<br>GAGAAGVLPG VGGAGVPGVP GAIPGIGGIA GVGTPAAAAA AAAAAKAAKY GAAAGLVPGG PGFGPGVVGV PGAGVPGVGV<br>PGAGIPVVPG AGIPGAAVPG VVSPEAAAKA AAKAAKYGAR PGVGVGGIPT YGVGAGGFPG FGVGVGGIPG VAGVPGVGGV<br>PGVGGVPGVG ISPEAQAAAA AKAAKYGVGT PAAAAAKAAA KAAQFGLVPG VGVAPGVGVA PGVGVAPGVG LAPGVGVAPG<br>VGVAPGVGVA PGIGPGGVAA AAKSAAKVAA KAQLRAAAGL GAGIPGLGVG VGVPGLGVGA GVPGLGVGAG VPGFGAVPGA<br>LAAAKAAKYG AAVPGVLGGL GALGGVGIPG GVVGAGPAAA AAAAKAAAKA AQFGLVGAAG LGGLGVGGLG VPGVGGLGGI<br>PPAAAAKAAK YGAAGLGGVL GGAGQFPLGG VAARPGFGLS PIFPGGACLG KAC |
| ELP29-36 | AAAAAKAAKY GAGVPGVGVP GVGVPGVGVP GVGVPGVGVP GVGVPGVGAG PAAAAAAAKA AAKAAQFGLV GAAGLGGLGV<br>GGLGVPGVGG LGGIPPAAAA KAAKYGAAGL GGVLGGAGQF PLGGVAARPG FGLSPIFPGG ACLGKACGRK RK |
| ELP29-36<br>ΔRKRK | AAAAAKAAKY GAGVPGVGVP GVGVPGVGVP GVGVPGVGVP GVGVPGVGAG PAAAAAAAKA AAKAAQFGLV GAAGLGGLGV<br>GGLGVPGVGG LGGIPPAAAA KAAKYGAAGL GGVLGGAGQF PLGGVAARPG FGLSPIFPGG ACLGKAC |
| ELP29-36<br>GD | AAAAAKAAKY GAGVPGVGVP GVGVPGVGVP GVGVPGVGVP GVGVPGVGAG PAAAAAAAKA AAKAAQFGLV GAAGLGGLGV<br>GGLGVPGVGG LGGIPPAAAA KAAKYGAAGL GGVLGGAGQF PLGGVAARPG FGLSPIFP**D**G ACLGKACGRK RK |
| ELP29-36<br>QSQS | AAAAAKAAKY GAGVPGVGVP GVGVPGVGVP GVGVPGVGVP GVGVPGVGAG PAAAAAAAKA AAKAAQFGLV GAAGLGGLGV<br>GGLGVPGVGG LGGIPPAAAA KAAKYGAAGL GGVLGGAGQF PLGGVAARPG FGLSPIFPGG ACLGKACG**QS QS** |
| ELP29-36<br>KA | AAAAAKAAKY GAGVPGVGVP GVGVPGVGVP GVGVPGVGVP GVGVPGVGAG PAAAAAAAKA AAKAAQFGLV GAAGLGGLGV<br>GGLGVPGVGG LGGIPPAAAA KAAKYGAAGL GGVLGGAGQF PLGGVAARPG FGLSPIFPGG ACLG**A**ACGRK RK |
| ELP$_3$ | AAAAAKAAKY GAGVPGVGVP GVGVPGVGVP GVGVPGVGVP GVGVPGVAAA AAKAAKYGAG VPGVGVPGVG VPGVGVPGVG<br>VPGVGVPGVG VPGVAAAAAK AAKYGAGVPG VGVPGVGVPG VGVPGVGVPG VGVPGVGVPG VAAAAAKAAK YGA |
| ELP$_3$(36) | AAAAAKAAKY GAGVPGVGVP GVGVPGVGVP GVGVPGVGVP GVGVPGVAAA AAKAAKYGAG VPGVGVPGVG VPGVGVPGVG<br>VPGVGVPGVG VPGVAAAAAK AAKYGAGVPG VGVPGVGVPG VGVPGVGVPG VGVPGVGVPG GACLGKACGR KRK |
| ELP$_3$(36)<br>ΔRKRK | AAAAAKAAKY GAGVPGVGVP GVGVPGVGVP GVGVPGVGVP GVGVPGVAAA AAKAAKYGAG VPGVGVPGVG VPGVGVPGVG<br>VPGVGVPGVG VPGVAAAAAK AAKYGAGVPG VGVPGVGVPG VGVPGVGVPG VGVPGVGVPG GACLGKAC |

**SI Table S2. Amino acid sequences of protein constructs used in this study.**

### Supplementary Figures

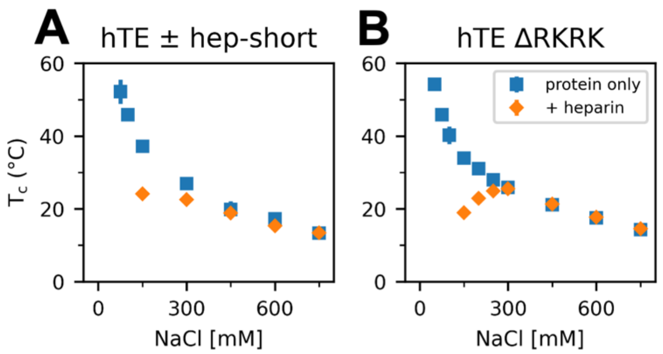


**SI Figure S1. Coacervation temperatures of hTE ± short heparin and hTE ∆RKRK ± heparin.** Coacervation temperatures for **(A) 25 µM** hTE ± 17.8 µM hep-short (short heparin, 6.3 kDa, fully sulfated) and **(B)** 25 µM hTE ∆RKRK ± 6.25 µM heparin (18 kDa, fully sulfated).

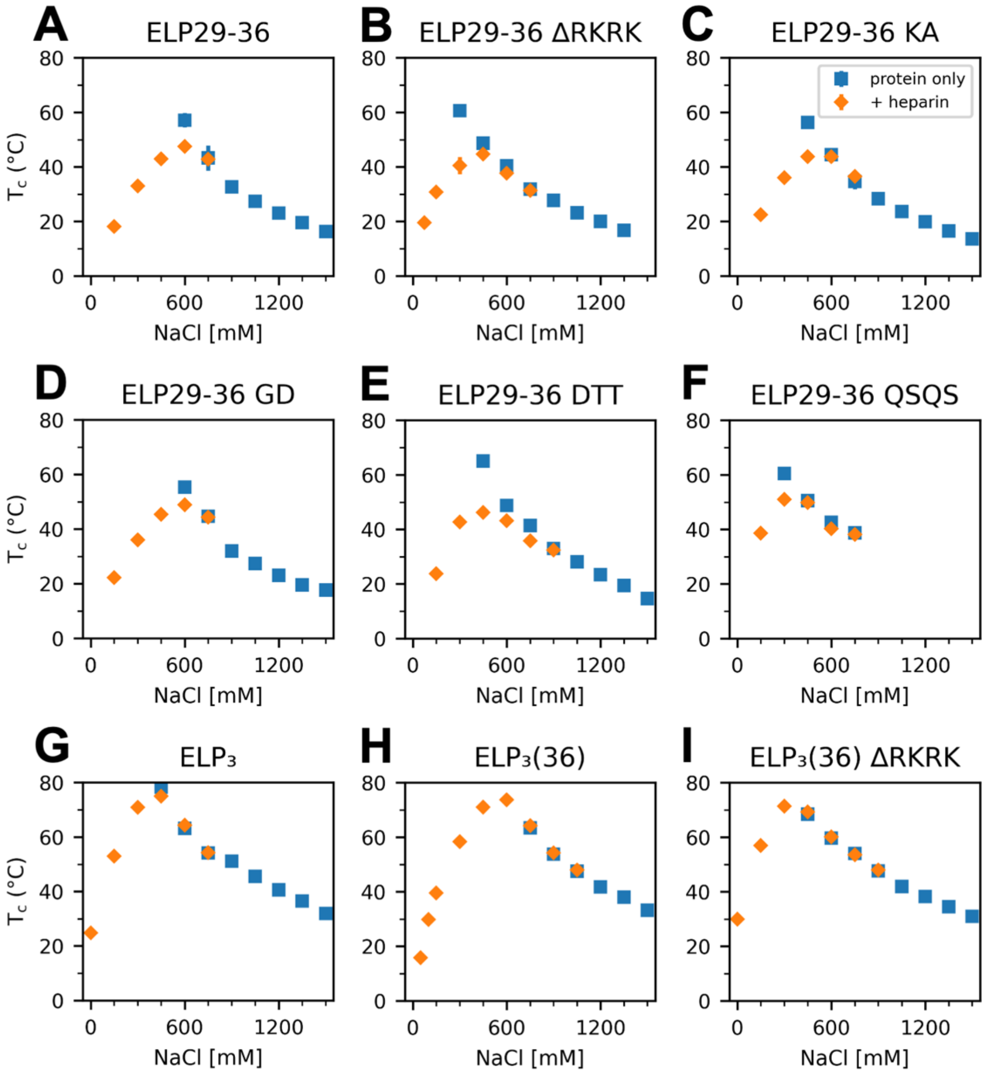


**SI Figure S2. Coacervation temperatures of ELP constructs containing domain 36 sequence variants.** Coacervation temperatures of ELP ± heparin for **(A)** ELP29-36, **(B)** ELP29-36 ΔRKRK, **(C)** ELP29-36 KA, **(D)** ELP29-36 GD, **(E)** ELP29-36 DTT, **(F)** ELP29-36 QSQS, **(G)** $ELP_3$, **(H)** $ELP_3(36)$, and **(I)** $ELP_3(36)$ ΔRKRK. Each sample contained 200 μM protein ± 50 μM heparin.

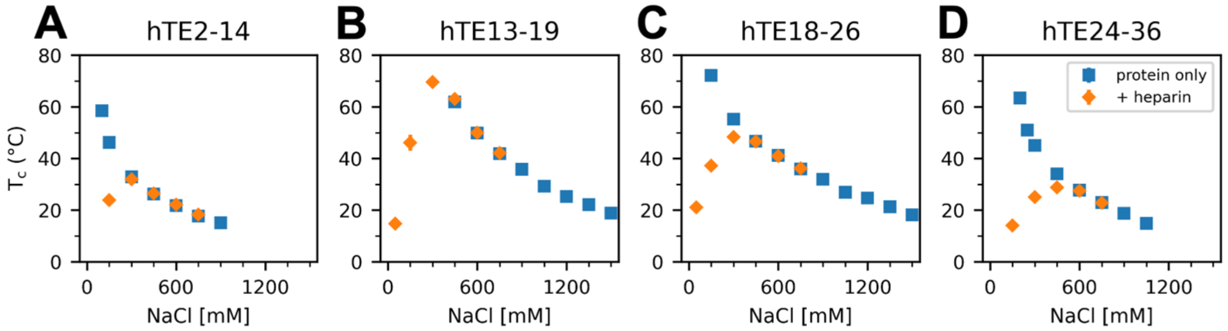


**SI Figure S3. Coacervation temperatures of tropoelastin fragments.** Coacervation temperatures for tropoelastin fragment constructs ± heparin for **(A)** hTE2-14, **(B)** hTE13-19, **(C)** hTE18-26, and **(D)** hTE24-36. Each sample contained 200 μM protein ± 50 μM heparin.

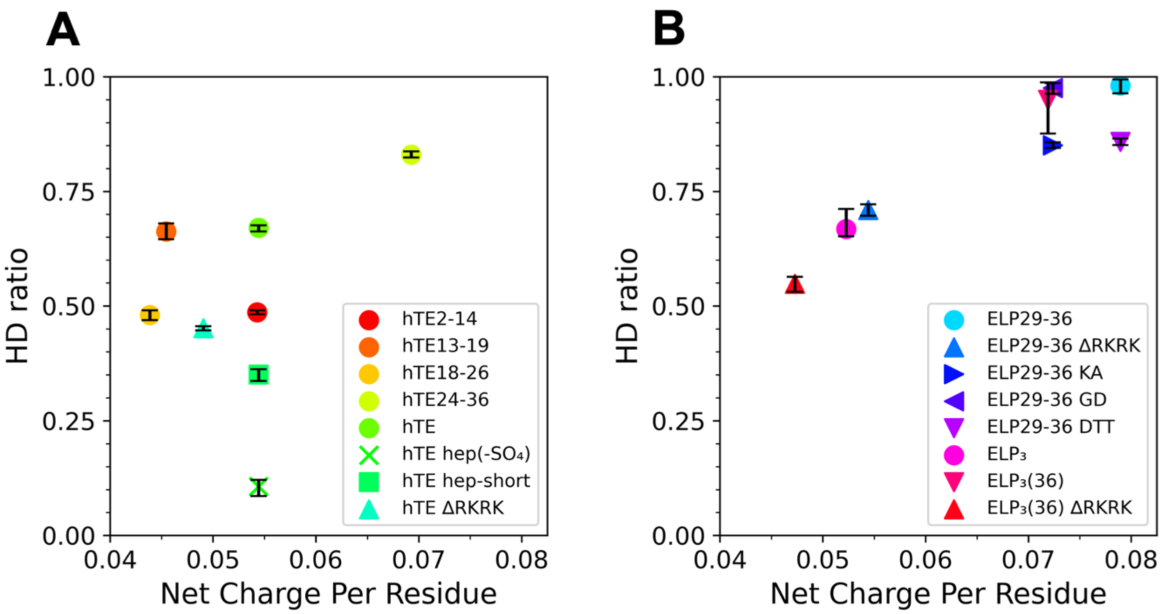


**SI Figure S4. Plots of HD ratio against sequence net charge per residue.** The heparin difference ratio (HD ratio) is shown as a function of net charge per residue (NCPR) for **(A)** tropoelastin and tropoelastin fragment constructs and **(B)** ELPs. Error bars are the standard deviation of the HD ratio.

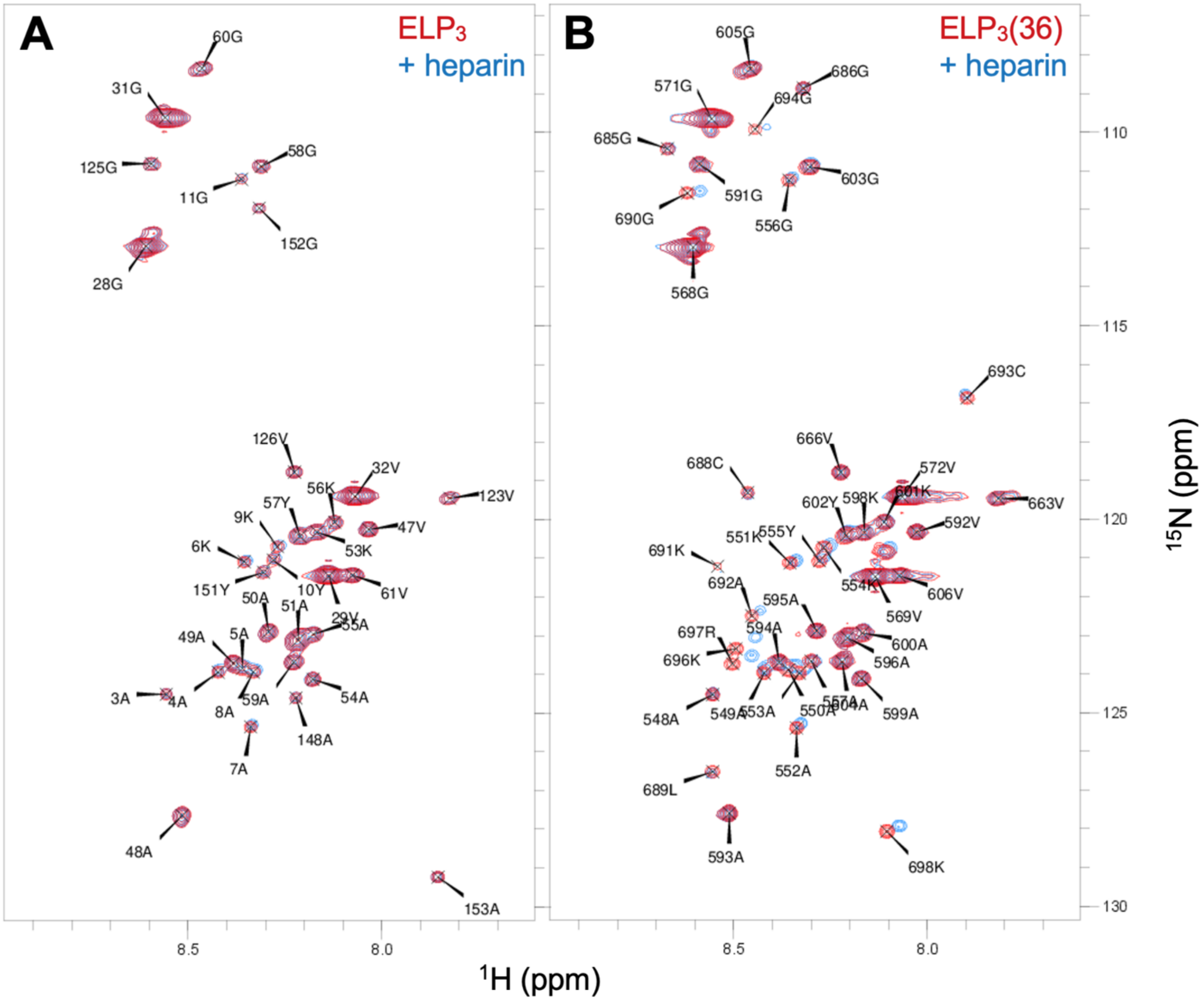


**SI Figure S5. Backbone $^1$H-$^{15}$N HSQC spectra of ELP$_3$ and ELP$_3$(36) in the presence and absence of heparin.** $^1$H-$^{15}$N amide backbone HSQC spectra of **(A)** ELP$_3$ and **(B)** ELP$_3$(36) at 200 μM without heparin (red) or in the presence of 50 μM heparin (blue). ELP$_3$ spectra were recorded at 600 MHz and ELP$_3$(36) spectra were recorded at 700 MHz. Buffers contain 100 mM NaCl, 50 mM sodium phosphate pH 7.2, 10% $D_2O$ and all data were acquired at 10 °C under non-coacervating conditions. The labels indicate assignments of well resolved peaks, with numbering of ELP$_3$ starting from 1, and numbering of ELP$_3$(36) such that the last residue of domain 36 shares nomenclature with the last residue of full-length tropoelastin (Lys 698).

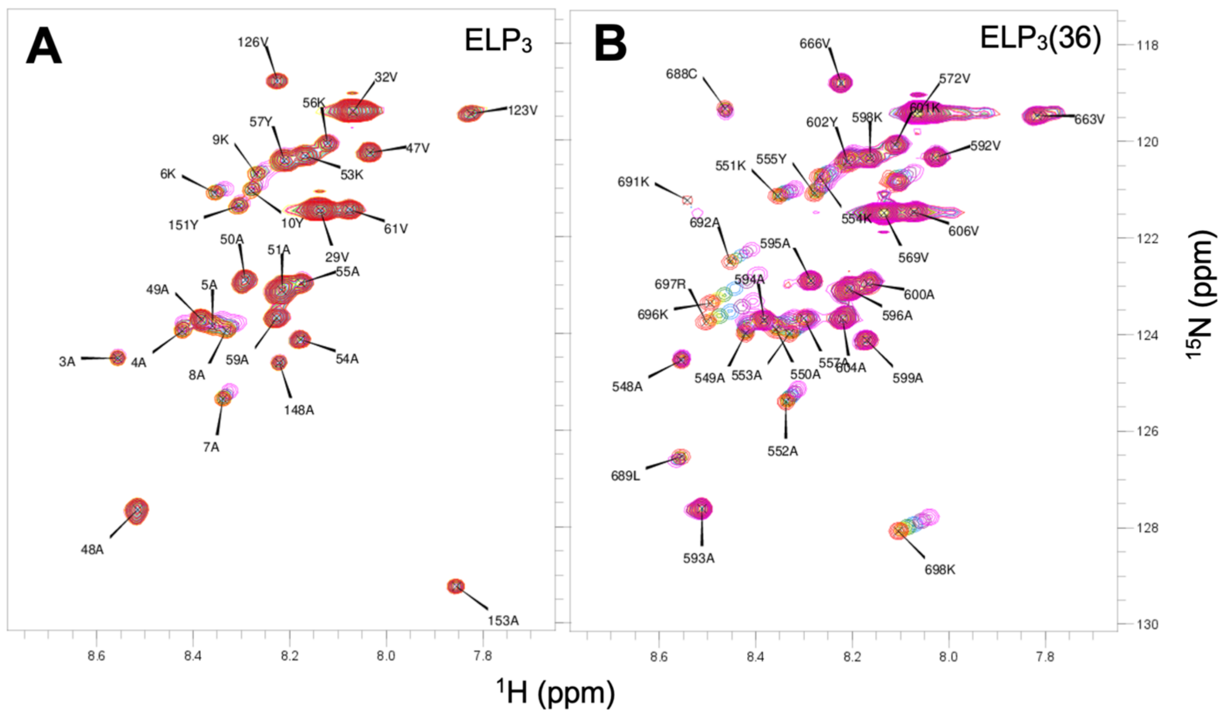


**SI Figure S6. Backbone $^{1}H$-$^{15}N$ HSQC upon titration of heparin into samples containing $ELP_3$ and $ELP_3(36)$.** $^{15}N$-$^{1}H$ HSQC spectra of samples containing 200 μM ELP and heparin concentrations from 0–200 μM (0:1–1:1 heparin:ELP molar ratio) are shown for **(A)** $ELP_3$ and **(B)** $ELP_3(36)$. Peak colours range from red (no heparin) to magenta (1:1 heparin:ELP molar ratio). Buffers contain 100 mM NaCl, 50 mM sodium phosphate pH 7.2, and 0.2% dioxane. $ELP_3$ spectra were recorded at 600 MHz and $ELP_3(36)$ spectra were recorded at 700 MHz. All data were obtained at 10 °C under non-coacervating conditions.

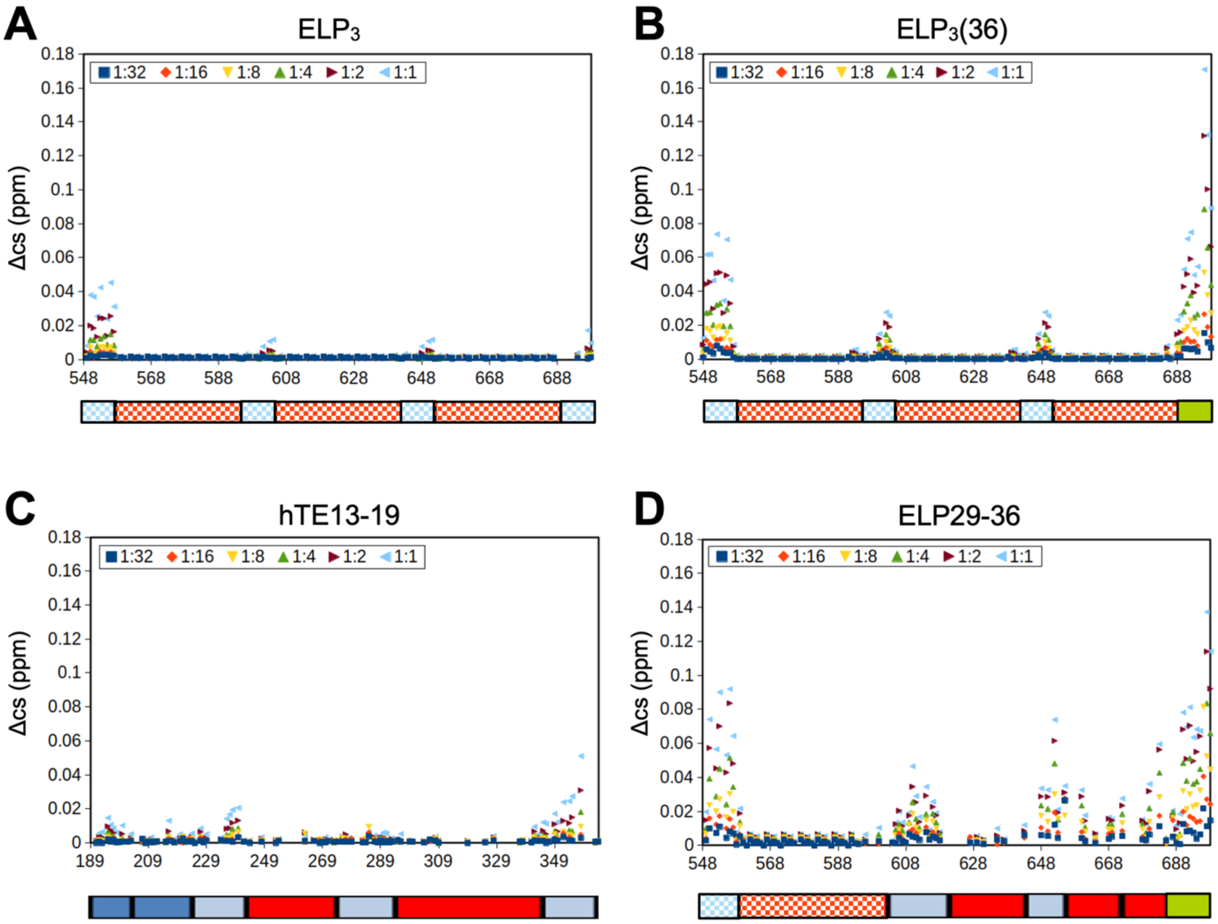


**SI Figure S7. Backbone $^{1}H$-$^{15}N$ chemical shift changes upon heparin titration.** The change in weighted backbone amide chemical shift (Eq. 3) as a function of increasing heparin concentration is shown for **(A)** $ELP_3$, **(B)** $ELP_3(36)$, **(C)** hTE13-19, and **(D)** ELP29-36. The legend shows the heparin:ELP molar ratios used, and the X-axis shows amino acid position within each construct. Schematics below each plot show construct domain architecture, as used in Figure 1 (red – hydrophobic domain, blue – crosslinking domain, green – domain 36, shaded – ELP domain).

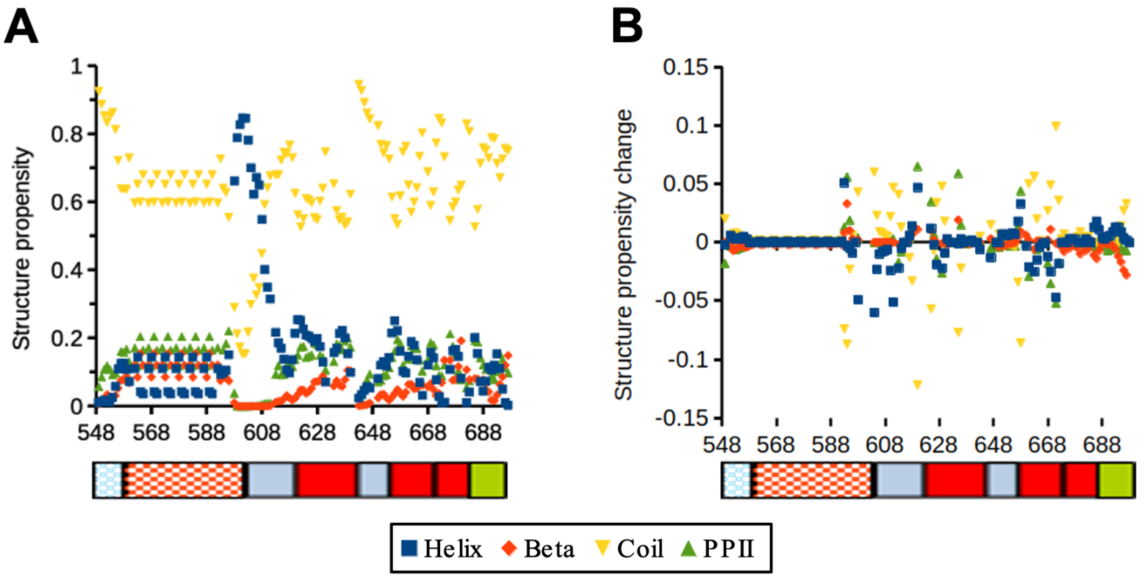


**SI Figure S8. Predicted secondary structure propensity for ELP29-36 in the presence of heparin. (A)** The secondary structure propensity predicted from backbone amide proton, nitrogen, and carbonyl carbon chemical shifts for 200 μM ELP29-36 in the presence of 25 μM heparin. **(B)** Change in structure propensity for ELP29-36 with and without heparin. Buffers contain 100 mM NaCl, 50 mM sodium phosphate pH 7.2, and NMR data were obtained at 700 MHz under non-coacervating conditions.

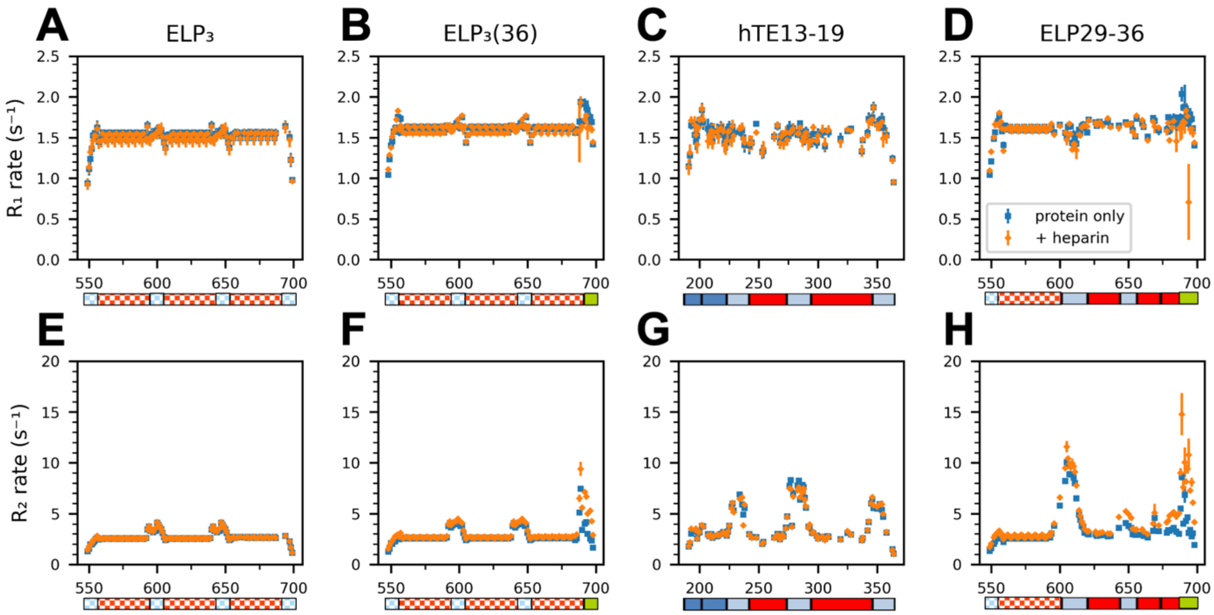


**SI Figure S9. $^1$H-$^{15}$N amide $R_1$ and $R_2$ relaxation rates of ELPs ± heparin.** $^{15}$N NMR relaxation rates obtained shown for 200 µM ELPs under non-coacervating conditions in the absence (blue) and presence of 50 µM heparin (orange). **[Top row]** $R_1$ relaxation rates: **(A)** $ELP_3$, **(B)** $ELP_3$(36), **(C)** hTE13-19, **(D)** ELP29-36. **[Bottom row]** $R_2$ relaxation rate: **(E)** $ELP_3$, **(F)** $ELP_3$(36), **(G)** hTE13-19, **(H)** ELP29-36.

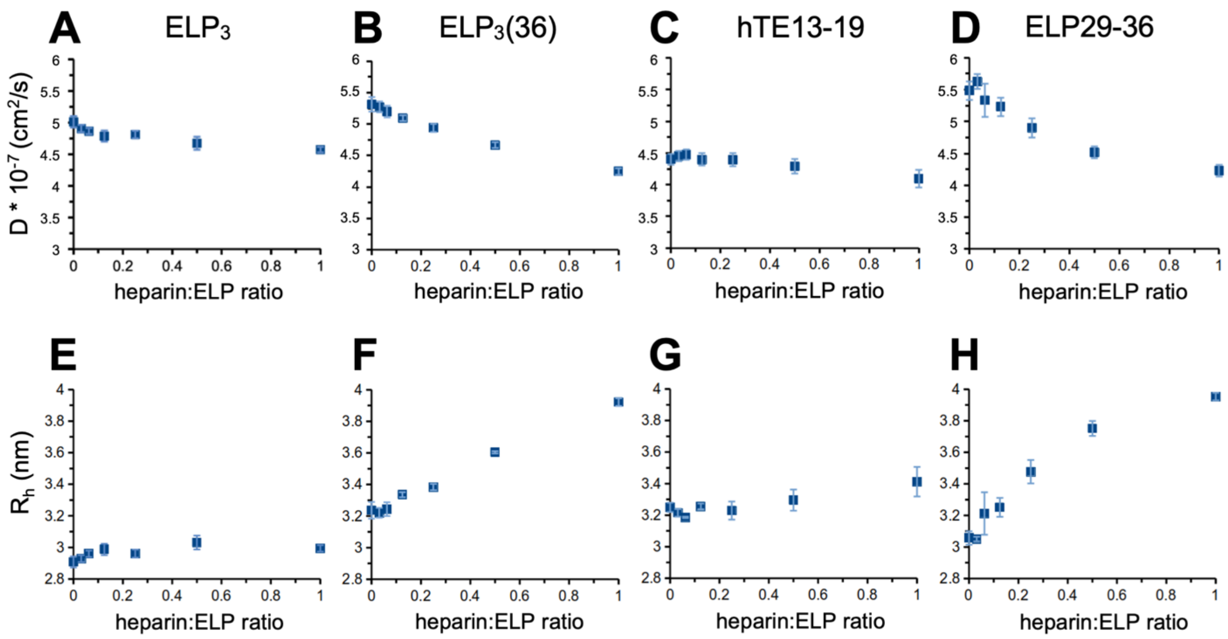


**SI Figure S10. Change in diffusion rate and estimated $R_h$ of ELPs upon heparin titration.** The hydrodynamic radii ($R_h$) of ELPs were estimated from NMR PFG diffusion measurements of the diffusion rate (D), with increasing heparin concentration from 0 – 200 μM (ELP concentration 200 μM, 0:1–1:1 heparin:ELP molar ratio). **[top row]** Diffusion rate, **[bottom row]** hydrodynamic radius for **(A, E)** $ELP_3$, **(B, F)** $ELP_3$(36), **(C, G)** hTE13-19, **(D, H)** ELP29-36. Error bars are the standard deviation of the fit error. Buffers contain 100 mM NaCl, 50 mM sodium phosphate, pH 7.2, and 0.2% dioxane. NMR diffusion measurements were performed at 600 MHz at 10 °C under non-coacervating conditions.

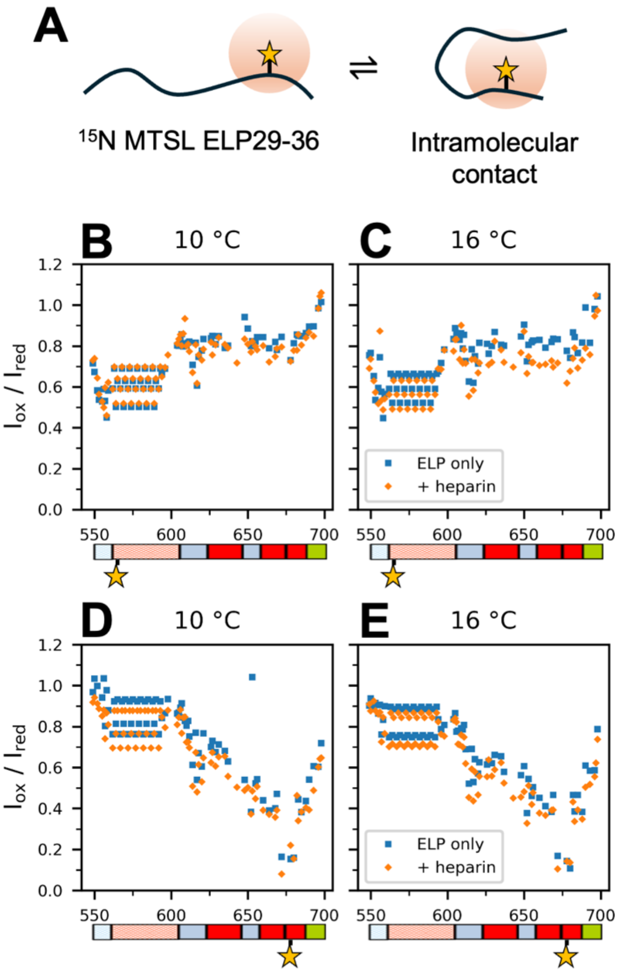


**SI Figure S11. Intramolecular PREs measured for ELP29-36 in the presence or absence of heparin. (A)** Schematic of the intramolecular PRE effect. As distal regions of the peptide backbone come within range of the paramagnetic MTSL label, their relaxation is enhanced and a reduction in signal intensity (measured as $I_{ox}/I_{red}$) is observed. **[Top row]** $I_{ox}/I_{red}$ data obtained for $^{15}$N-labeled ELP29-36 with MTSL attached to an N-terminal cysteine, recorded at a temperature of **(B)** 10 °C or **(C)** 16 °C. **[Bottom row]** $I_{ox}/I_{red}$ data obtained for $^{15}$N-labelled ELP29-36 with MTSL attached to a C-terminal cysteine, recorded at **(D)** 10 °C and **(E)** 16 °C. Note that the coacervation temperature in the presence of heparin is 18 °C. The domain organization of ELP29-36 and location of the MTSL tag is shown below the horizontal axis for reference. All samples contained an equimolar amount of unlabeled protein to reduce intermolecular PRE effects and to maintain the same total ELP concentration and coacervation temperature as the intermolecular PRE samples (Figure 6).
